# Boundary constraints can determine pattern emergence

**DOI:** 10.1101/2025.07.21.665949

**Authors:** Yi Ting Loo, Juliet Chen, Ryan Harrison, Tiago Rito, Sophie Theis, Guillaume Charras, James Briscoe, Timothy E. Saunders

## Abstract

The robust patterning of cell fates during embryonic development requires precise coordination of signalling gradients within defined spatial constraints. Using a geometrically confined *in vitro* system derived from human embryonic stem cells, we demonstrate that patterning of neuromesodermal progenitors (NMPs) during axial elongation is driven by boundary-dependent mechanisms. Despite extensive work on radial fate patterning in confined 2D systems, the quantitative role of boundary conditions in shaping spatiotemporal dynamics remains unclear. Here, we show that a minimal reaction-diffusion model coupled with a simplified gene regulatory network accurately predicts spatial patterns across diverse geometries. Guided by its predictions, we identify Wnt signalling as a key component of the activator signal. Inhibition of Wnt secretion preserved initiation of patterning but disrupted subsequent morphogenesis, indicating distinct mechanisms govern pattern establishment versus maintenance. Our findings reveal how geometry encodes positional information that directs molecular patterning, providing insight into how spatial constraints and signalling dynamics guide robust tissue self-organisation during development.

## INTRODUCTION

Embryonic development relies on robust collective self-organisation to sculpt complex tissue architectures. Cells display a broad range of behaviours, including differentiation, proliferation and migration^1^. Such processes typically occur within spatial boundaries, which can be in the form of surrounding tissues, an external shell, or compartmentalised ECM. These boundaries can shape key aspects of development, including morphogen gradients, cell positioning, and force generation^2–6^.

The influence of boundaries on morphogenesis is pervasive, beginning soon after fertilisation. For example, in early *C. elegans* embryos, spatial confinement of blastomeres governs their interactions and signalling environment^7^, with physical constraints imposed by the shell essential for consistent and robust blastomere positioning^8,9^. Boundaries continue to influence development beyond early stages, guiding large-scale tissue organisation and body axis formation. In vertebrate embryos, the body axis progressively extends through a process of axial elongation, during which neural and mesodermal tissues are laid down in precise proportions and spatial domains along the anterior-posterior axis^10–13^. Volumetric growth during elongation is driven by multipotent axial progenitors in the tailbud, which continuously differentiate and exit the progenitor zone when committed to a fate. The posterior presomitic mesoderm (pPSM) exhibits intrinsic motility, believed to generate the forces required for elongation; however, directional tissue extension only arises in the presence of physical boundaries^14,15^ as observed in *ex vivo* avian pPSM explants. This demonstrates that spatial constraints are essential for guiding morphogenesis^16,17^.

Due to the difficulty of isolating cellular behaviour, tissue mechanics, and confinement effects *in vivo*, the role of boundaries in vertebrate patterning and axial elongation remains poorly understood. To address this challenge, several *in vitro* 2D organoid models derived from pluripotent stem cells (PSCs) have been developed, using substrate micropatterning to impose spatial constraints. These provide tractable, reproducible systems to study developmental mechanisms (reviewed in^18^). Systems such as 2D gastruloids have been used extensively to study cell fate specification into germ layers^19–21^. Under confinement, these gastruloids generate robust concentric radial patterns of ectoderm, mesoderm and endoderm^13,21,22^.

Building on gastruloid models, posterior neuruloids were developed to specifically capture trunk elongation^23^. Here, cycles of differentiation on circular micropatterns produce radially organised cell fates, with mesodermal cells forming near the colony edge and neural cells toward the centre. By day 3, neuruloids undergo a morphogenetic transition into a complex 3D structure, hypothesised to be driven by signals originating at the colony boundaries that organise and guide axial structure formation. Transcriptomic profiling of day-3 neuruloids confirmed their posterior identity, characteristic of mid-to late-stage elongation, indicating that neuruloid development encompasses bipotent neuromesodermal progenitors (NMPs) and their downstream mesodermal and neural lineages.

Under confinement, the first symmetry-breaking event after neuruloid induction is the emergence of a peripheral ring of cells expressing the primitive streak and mesodermal marker Brachyury (TBXT). The rapid emergence of radial patterning is a shared feature of micropatterned developmental models, although the proposed underlying mechanisms vary between systems. In BMP-induced gastruloids, patterning has been attributed to the relocalisation of BMP receptors, which become apically exposed only at the colony edge, restricting morphogen signalling to peripheral cells^24^. In Wnt-treated colonies, TBXT expression initially appears throughout the colony before becoming confined to the edge, driven by E-Cadherin-dependent cell density differences^22^. Reduced E-Cadherin junctions at the periphery increase cytoplasmic *β*-catenin and Wnt sensitivity, while inhibitors accumulate toward the centre.

In neuruloids, rather than using Wnt ligand, Wnt signalling is activated using Chiron (CHIR), a small molecule that acts cell-autonomously without involving cell surface receptors or secreted inhibitors. TBXT patterning in neuruloids shows gradual edge restriction by 24 hours, whereas previously on micropatterned hESCs, CHIR treatment abolished all spatial TBXT organisation^22,23^. This indicates that posterior fate patterning arises through distinct mechanisms and highlights an important role for spatial constraints in defining initial cell fate domains. However, how these patterns respond to changes in boundary conditions, and how robust boundary-driven patterning is across length and time scales has not yet been thoroughly quantified.

Alongside improvements in experimental models of tissue patterning, there have recently been significant advances in theoretical approaches to understand cell fate determination^25^ and robust patterning^26–28^. Reaction-diffusion theory, originally proposed by Alan Turing^29^, remains one of the most powerful and widely used frameworks for modelling spontaneous symmetry breaking and pattern formation in initially homogeneous systems. Alongside Wolpert’s concept of positional information^30,31^, such models have successfully captured key features of diverse biological processes^26,32,33^. Notably, these models are particularly well-suited for explaining phenomena where patterns emerge without pre-existing spatial cues. Despite differences in biological context and specific mechanisms of each system, reaction-diffusion models have been applied to study the formation of biological patterns across a range of spatial scales, from subcellular^34^, to tissue^28,35,36^, and up to population dynamics in ecological systems scales^37^. Importantly, this approach can make quantitative predictions about the system’s behaviour without requiring precise molecular or mechanistic details^26,28,38,39^. However, it remains unclear how well a minimal reaction-diffusion model can account for spatially constrained patterning in conditions where symmetry is broken within defined geometric boundaries.

In this study, we show that NMP-driven morphogenesis is sensitively and predictably guided by boundary constraints. Two stages of symmetry-breaking events occur during neuruloid development: the emergence of TBXT radial patterning at day 1 post-induction and subsequent TBXT cluster formation by 3 days post-induction. Both events can be predicted using reaction-diffusion models. We show that TBXT pattern emergence is consistent with induction by a diffusive boundary-derived signal (*e*.*g*. a morphogen) acting over a defined length scale^30,40^. When coupled to a minimal gene regulatory network (GRN), the model predicts how boundaries affect the balance between competing cell fates marked by the NMP genes SOX2 and TBXT. Overall our findings demonstrate that boundary constraints can provide positional information through reaction-diffusion principles, offering insight into the architecture of the gene regulatory network that underlies spatial pattern emergence.

## RESULTS

### A phenomenological reaction-diffusion model robustly captures neuruloid patterning

To investigate initial pattern emergence during vertebrate trunk formation, we used *neuruloids*, an organoid model of posterior trunk development^23^. Human embryonic stem cells were seeded onto circular, laminin-coated micropatterns to impose spatial confinement. Starting from a homogeneous population, neuruloids self-organised over the course of 3 days^21,23,41^. Specifically, the initially uniform layer of pluripotent SOX2-expressing cells differentiated following induction with Wnt/FGF agonists and BMP/NODAL inhibitors (Fig. 1A), leading to the emergence of neuromesodermal progenitors (NMPs) - precursors to posterior neural and trunk mesodermal lineages^23^. During its development, the monolayer underwent morphological changes as a second layer of cells gradually emerged at the neuruloid periphery, transitioning the structure into 3D (Fig. 1B, lower panel), consistent with previous observations. To analyse the patterning and morphogenetic dynamics, neuruloids were fixed at defined time points - 6, 24, and 48 hours postinduction. Prior to this morphogenetic transition, we confirmed a robust initial symmetry-breaking event in which TBXT expression became restricted to a peripheral ring, marking mesodermal differentiation. (Fig. 1A-B)^23^. *In vivo*, the presomitic mesoderm drives mechanical elongation^14^. Hence, this early pattern of mesodermal induction, which appears to be boundary-localised, may contribute to initiating the morphogenetic changes that follow.

**Figure 1:**
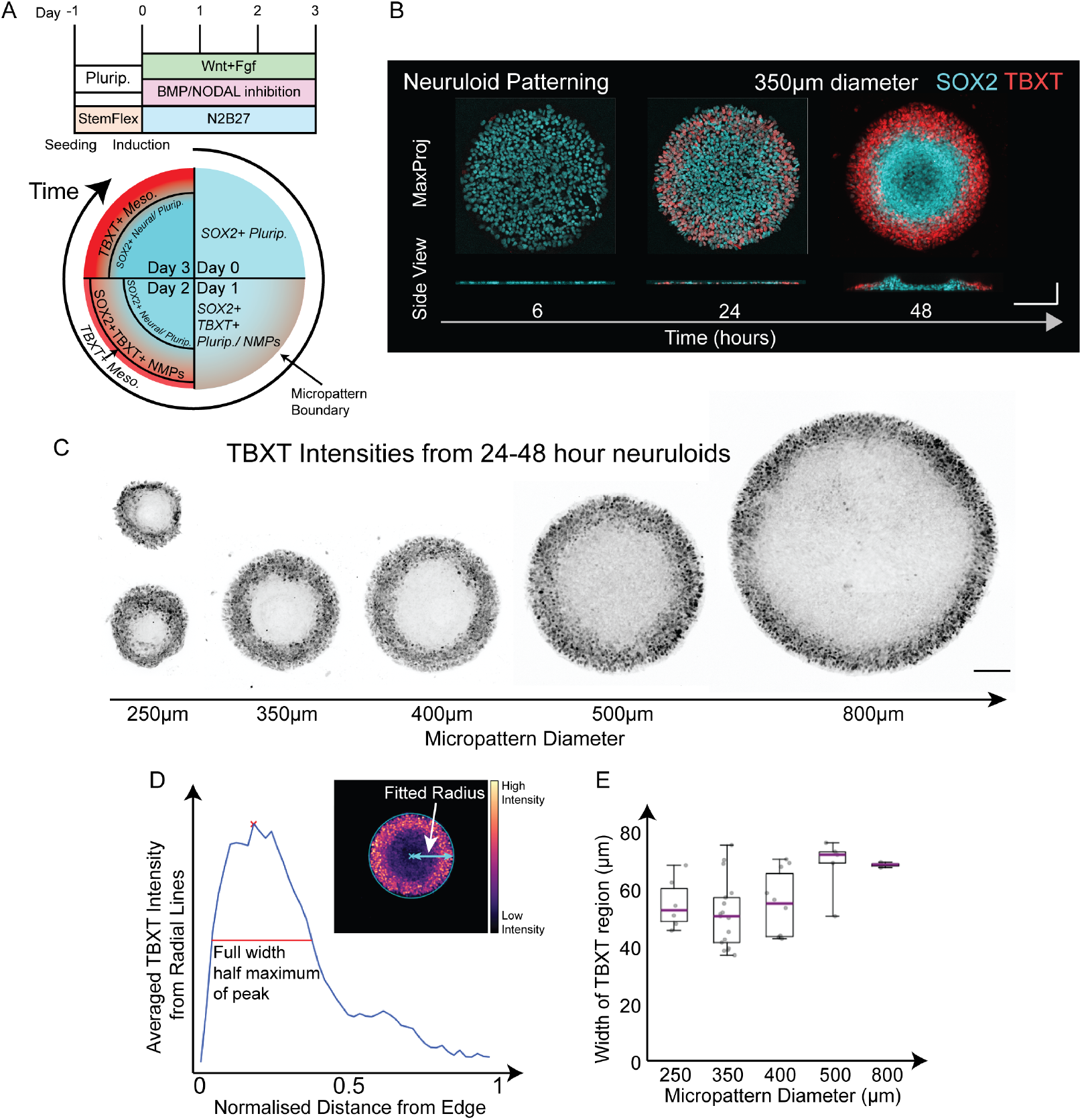
Neuruloids reproducibly pattern SOX2 and TBXT positive cells upon induction. **A**. Schematics of neuruloid protocol and cell fate patterning during neuruloid development, specifically pluripotent, neuromesodermal progenitors (NMPs), neural and mesoderm cells, upon induction. **B**. Patterning timeline during the first two days of neuruloid development post-induction. Maximum projections show radial patterning of TBXT expression, which was reproducible across multiple experiments. Crosssectional views show an initially homogeneous layer of cells developing to a 3D shape at 48 hour postinduction. **C**. TBXT patterning in neuruloids grown in circular micropatterns of different sizes. **D**. To estimate the widths of TBXT ring, we obtain the full width half maximum (FWHM) of the peaks from the TBXT intensity plots. **E**. Quantification of the widths of TBXT ring, where *n* = 6, 17, 10, 5, 2 neuruloids for sizes 250, 350, 400, 500 and 800*µ*m respectively were used for quantification. The boxplots display the interquartile range (IQR), with a line indicating the median. The whiskers extend from the box to the most extreme data points lying within 1.5x the (IQR). No statistically significant difference was found between the 5 groups of sizes (Kruskal-Wallis test, H=9.085, p=0.059). Scale bars: All horizontal scale bars are 100*µ*m, vertical scale bar is 50*µ*m.

To investigate the early radial pattern emergence of TBXT expression, we test the hypothesis that TBXT expression is induced by proximity of the cells to the boundary. In particular, we reasoned that the width of this region should be invariant to changes in colony size. We examined this by varying the diameter of the micropatterns and analysing the resulting cell fate patterning. We found that, despite substantial size differences, a radial pattern consistently emerged 24 hours post-induction at the neuruloid periphery (Fig. 1C). By quantifying the width of the TBXT-positive ring in 24 to 48-hour neuruloids, we found that the spatial extent of TBXT expression pattern was invariant to changes in system size (Fig. 1D-E and S1A-B); the width of the TBXT ring remained consistent, with no significant differences between colony sizes (Fig. S1B). These observations suggest that TBXT patterning occurs within a characteristic distance from the boundary, consistent with a morphogen gradient of fixed range driving pattern emergence. Therefore, we hypothesised that the spatial extent of TBXT expression is determined by how cells respond to signalling molecules in relation to the boundary, which can act as either a source or a sink of morphogen.

To gain theoretical insight into how such a mechanism patterns colonies, we considered two phenomenological models of early radial TBXT patterning based on diffusing molecules (Fig. 2A). The simplest model is a diffusion–degradation model for a single diffusible activator of TBXT (Supplementary Sec. S2), whereas a more complex reaction–diffusion model consists of two coupled partial differential equations (PDEs) capturing the interaction between a diffusible activator of TBXT and its endogenous inhibitor (Fig. S1C) (Supplementary sec S3). A key question is how to best encode the observed effects in the boundary conditions? In micropatterned colonies, edge sensing has previously been attributed to boundary-restricted receptor accessibility to ligands in the culture medium^21,22,42^ (Fig. S1C). However, in our system the primary activator is Chiron (CHIR), a Wnt agonist that acts independently of cell surface receptors, yet neuruloids still exhibit clear edge sensing. To model edge detection, we applied a mixed boundary condition^43^: to both models, we applied a fixed activator concentration to impose a uniform state at the boundary, representing constant cellular sensitivity to signalling, and for the second activator-inhibitor model, a no-flux condition for the inhibitor is applied to reflect its endogenous secretion (Fig. S1C). Fixing the activator concentration at the boundary for reaction-diffusion models isolates the pattern formation to the interior of the domain. Enforcing the role of a consistent boundary reduces sensitivity to initial perturbations, thus promoting robust and biologically meaningful patterns^43^. The equations were solved numerically on a two-dimensional domain using the finite element method, yielding the predicted steady-state activator concentration profile across the domain.

**Figure 2:**
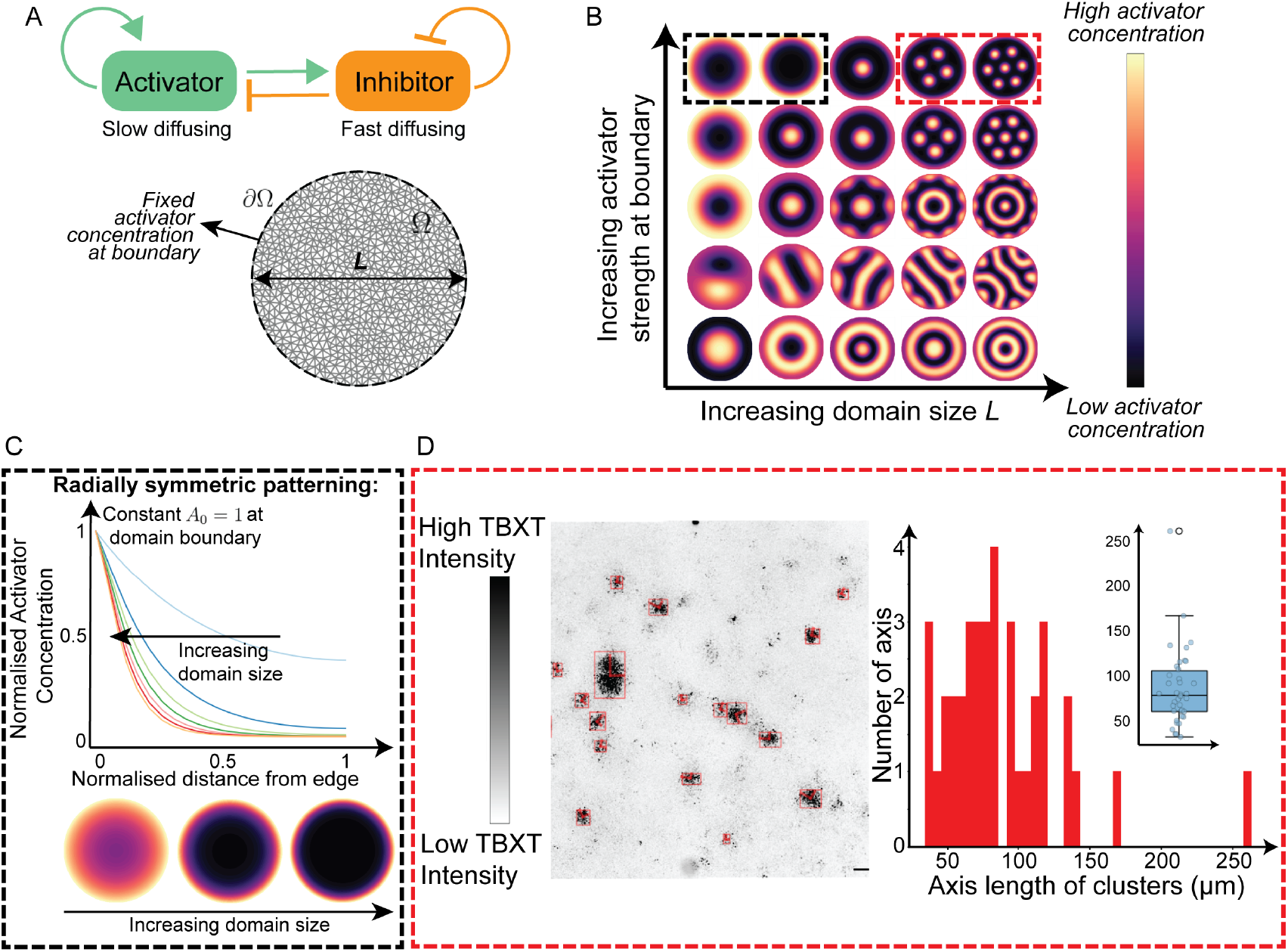
Reaction-diffusion modelling as a phenomenological model for TBXT patterning in neuruloids. **A**. Schematic of activator-inhibitor interaction modelled using a reaction-diffusion model (Supplementary Material, S3 for details). **B**. Parameter sensitivity analysis of the reaction-diffusion model, with varying domain length and strength of activator at the boundary. Black and red dashed boxes are referred to in **C** and **D** respectively. **C**. Relative activator concentrations across increasing domain sizes on a normalised spatial axis show radial pattering with increasing gradient steepness. **D**. (Left) Neuruloids grown on an unconstrained domain show clusters of high TBXT region. (Right) Quantification of the axis lengths of the clusters as a histogram (red bars) and corresponding boxplot (blue) showing the distribution. The boxplot displays the interquartile range (IQR), with a line indicating the median. The whiskers extend from the box to the most extreme data points lying within 1.5x the (IQR).

For the first simple diffusion-degradation model, we were able to phenomenologically reproduce the radially symmetric patterning seen from TBXT expression data for increasing micropattern sizes (Fig. S2 and Supplementary Sec. S2). For the reaction-diffusion model consisting of an activator-inhibitor pair, we performed sensitivity analysis to explore the patterns that can be obtained from changing the domain size and strength of the fixed activator concentration at the boundary relative to concentrations at equilibrium in the absence of diffusion (Fig. 2B, S1D and Supplementary sec S3). We focused on two regions, highlighted by dashed boxes in Fig. 2B. In the first region (black box), we identified parameter sets that produced radially symmetric patterns resembling the TBXT expression observed in neuruloids. This pattern is inverted when boundary input is lower than the concentration at equilibrium, concentrating high activator levels at the centre rather than the boundary. As the domain size increased in the numerical simulations, the activator concentration profile on a normalised spatial domain steepened, similar to the results from the first model (Fig. 2C and Fig. S2B). This aligns with the earlier observation that TBXT-positive regions maintain a consistent width, regardless of the overall size of the micropattern. As a result, in larger micropatterns, TBXT expression appears more restricted relative to the total area (Fig. 1C and S1A).

Other parameter sets led to a broad variety of patterns, including stripes and concentric rings (Fig. 2B and Supplementary sec S3 for details). Further increasing the domain size in the model led to the appearance of multiple foci of high activator concentration, rather than a single boundary-localised domain (Fig. 2B, red dashed box). We experimentally tested this prediction by seeding cells at the same density on a much larger substrate (18 mm diameter). In agreement with the model, we observed the formation of spatially separated clusters of cells expressing high TBXT (Fig. 2D, left). In contrast, the simple diffusion–degradation model was unable to reproduce this (Fig. S2 and Supplementary Sec. S2), supporting the requirement for an activator–inhibitor pair in neuruloid patterning. Quantification of cluster sizes revealed that their diameters closely matched the characteristic width of TBXT radial patterning observed in circular micropatterns (Fig. 2D, right). In summary, our phenomenological reaction-diffusion model effectively captures the TBXT spatial patterning observed in neuruloids with a length scale set by the system boundary, and will be used as the model for the rest of this section.

### Predicted activator profiles align with geometry-dependent patterning

We then sought to test the predictive power of our reaction-diffusion framework. If boundary constraints determine the domain of TBXT expression, we reasoned that altering colony geometry should change the spatial range and organisation of the TBXT domain. We first parameterised our model using the circular domains as calibration data.

To do this, we quantified the TBXT signal across multiple circular domains, accounting for variations in cell density by normalising using DAPI signal (Fig. S3A-C). Since our reaction-diffusion system is a phenomenological model, we fixed parameters that qualitatively replicated the observed TBXT radial patterns through sensitivity analysis, such as the activator concentration at the boundary (Fig. S1D). This reduced the model to a single free parameter. We fitted the simulated activator profile to the experimental TBXT expression profile (Fig. 3A) to find the relation between the dimensionless length scale of the model, *γ*, and the actual micropattern domain size. More details on the dimensionless parameters of the model and fitting can be found in the Supplementary Material, Section S3. We kept all the parameters and the relation between the length scale, *γ*, and area of micropattern constant in our analysis below, signifying that our model has no free parameters.

**Figure 3:**
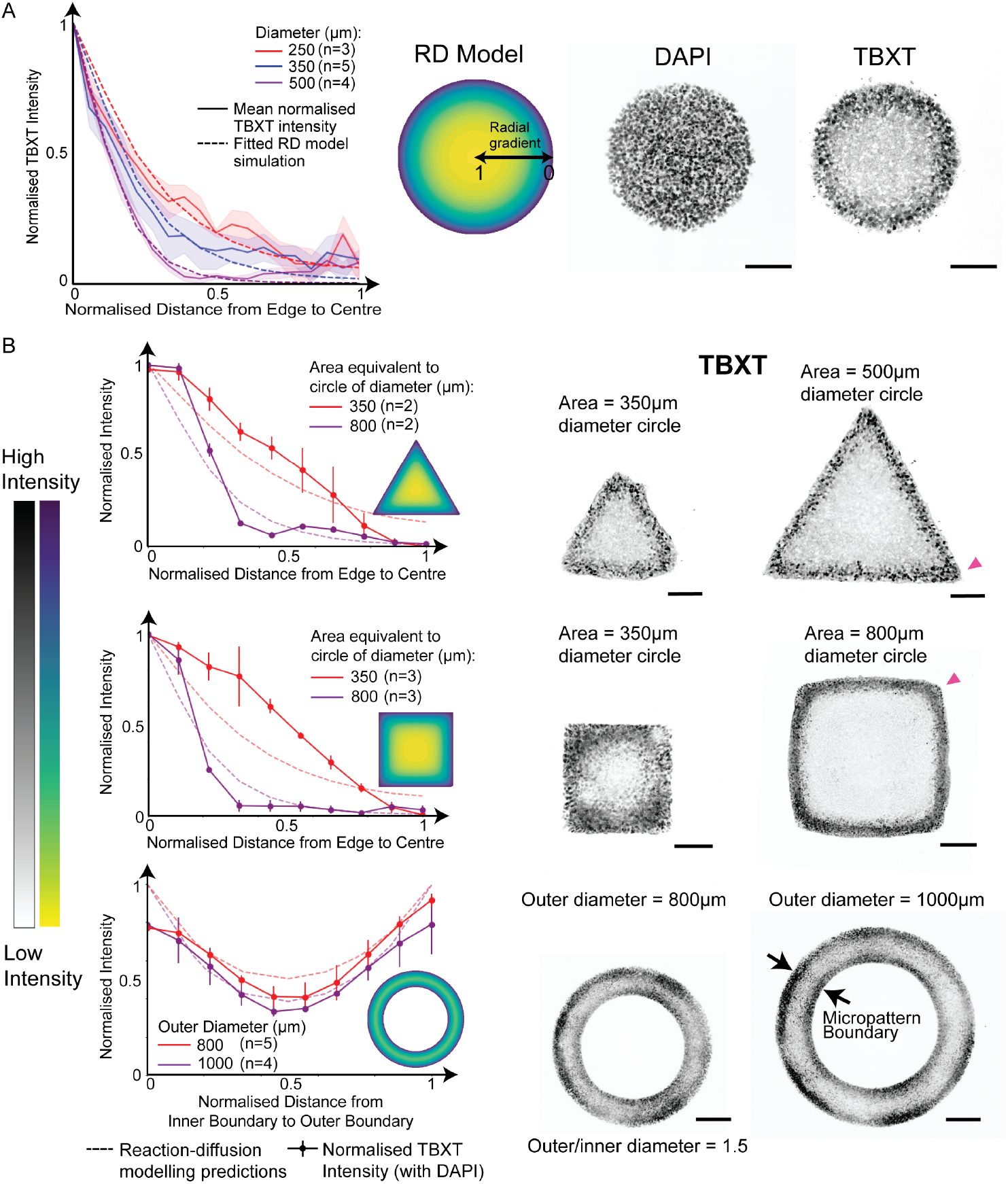
TBXT patterning in neuruloids of varying geometry. **A**. DAPI and TBXT intensities of circular neuruloids were used to extract normalised TBXT expression. The radial gradient generated by the reaction-diffusion model was fitted to the experimental TBXT expression profile (see also Fig. S3). (Right) Normalised TBXT expression. Solid lines and shaded regions show the mean and standard deviation respectively. At least *n* = 3 for each micropattern diameter. Dashed lines show the best fitted gradient from the model to the data. **B**. (Left) Neuruloids grown on micropatterns of different shapes. Micropattern sizes correspond to the areas of circular micropatterns of specific radii. The heatmap shows TBXT intensity. Magenta arrows indicate outer convex boundary (annulus) and corners (triangle and squares), cyan arrow indicates the inner concave boundary of annulus. (Right) Quantification of TBXT intensity in different geometries. Error bars show s.d. (see also Fig. S4B-C). Dashed lines are predictions from the reaction-diffusion model. Scale bars: All scale bars are 100*µ*m.

To investigate the influence of geometric confinement on pattern formation, we generated model predictions of activator intensity profiles on domains of various shapes: annuli, triangles, and squares (Fig. 3B first column (dashed curves and heatmap) and S4A). Triangle and square geometries allowed us to assess whether the presence of straight edges and sharp vertex angles influences the positioning of cell fate boundaries. Activator distributions on these domains were predicted to be edge-restricted, similar to circular domains. Annulus shapes feature both inner and outer boundaries, and activator levels were predicted to be high at both boundaries. This is as expected given the boundary conditions of the model.

To experimentally verify this, we grew neuruloids on these micropattern geometries (Fig. 3B, second and third columns). With the exception of annuli, all shapes were matched in area to the circular domains previously used. In general, pattern formation on these shapes displayed TBXT expression in bands of widths comparable to those on circular micropatterns (Fig. 3B and S4B-C). On edges, TBXT width was not different from the width on circular domains but we observed an expansion of the TBXT signal near corners. This effect was more pronounced on triangular domains than on squares, suggesting that sharper angles enhanced TBXT induction (Fig. S4B). When comparing experimental TBXT expression with predictions from the reaction–diffusion model, we observed strong agreement in the overall trend between their spatial profiles (Fig. 3B, first column). Notably on annuli, TBXT expression emerged at both boundaries, aligning with predictions, providing further evidence that edge-associated chemical cues drive self-organisation in this system. However, the experimental intensity profiles for both triangular and square micropatterns deviate from the model in a similar manner by comparing the solid and dashed lines in Fig. 3B: TBXT levels are higher in the mid-radius region for smaller areas but lower in the same region for larger areas. This indicates that certain key aspects of the underlying phenomenon are not fully captured by the model. It is important to note that the reaction–diffusion model describes molecular concentration fields rather than TBXT gene expression dynamics, even though a correlation between the two is assumed. Consequently, the correspondence in spatial trend, rather than absolute intensity, is the primary measure of agreement in this comparison. The observed shift of high TBXT signal intensity away from the edges in smaller micropatterns likely arises from mechanical influences or differences in the 3D morphological development of these neuruloids.

### A minimal gene regulatory network predicts curvature-dependent SOX2 and TBXT patterning

We have so far assumed that a signal emanating from the boundary activates TBXT expression. However, the underlying interactions between downstream transcriptional regulators are typically more complicated^25,44,45^. To investigate how activator concentration regulates the expression of these genes, we developed a minimal model with ordinary differential equations (ODEs) to capture the gene regulatory network (GRN) of SOX2 and TBXT. As we are principally interested in the initial symmetry breaking event, we focused only on the early interactions of genes of interest in the system and excluded contributions from other downstream genes.

The balance between SOX2 and TBXT expression is crucial for maintaining the progenitor state of NMPs. However, during lineage commitment, these transcription factors act antagonistically^46,47^. This mutual inhibition, together with local Wnt/FGF signalling levels, plays a key role in directing cell fate toward mesodermal (high TBXT) or neural (high SOX2) lineages^48^. Accordingly, in our ODE model, SOX2 and TBXT mutually repress one another (Fig. 4A), similar to the interactions used in previous systems^42^. We hypothesised that the activator stimulates TBXT expression. Thus, in this GRN, the activator concentration from the reaction-diffusion model (step 1) is a direct input to the ODE model (step 2), providing spatial information (Fig. 4A, Methods and Supplementary sec S5). Given that SOX2 is highly-expressed in self-renewing stem cells, we incorporated self-activation of SOX2 into the model to reflect its role in maintaining pluripotency and sustained expression in the absence of differentiation cues^49^.

**Figure 4:**
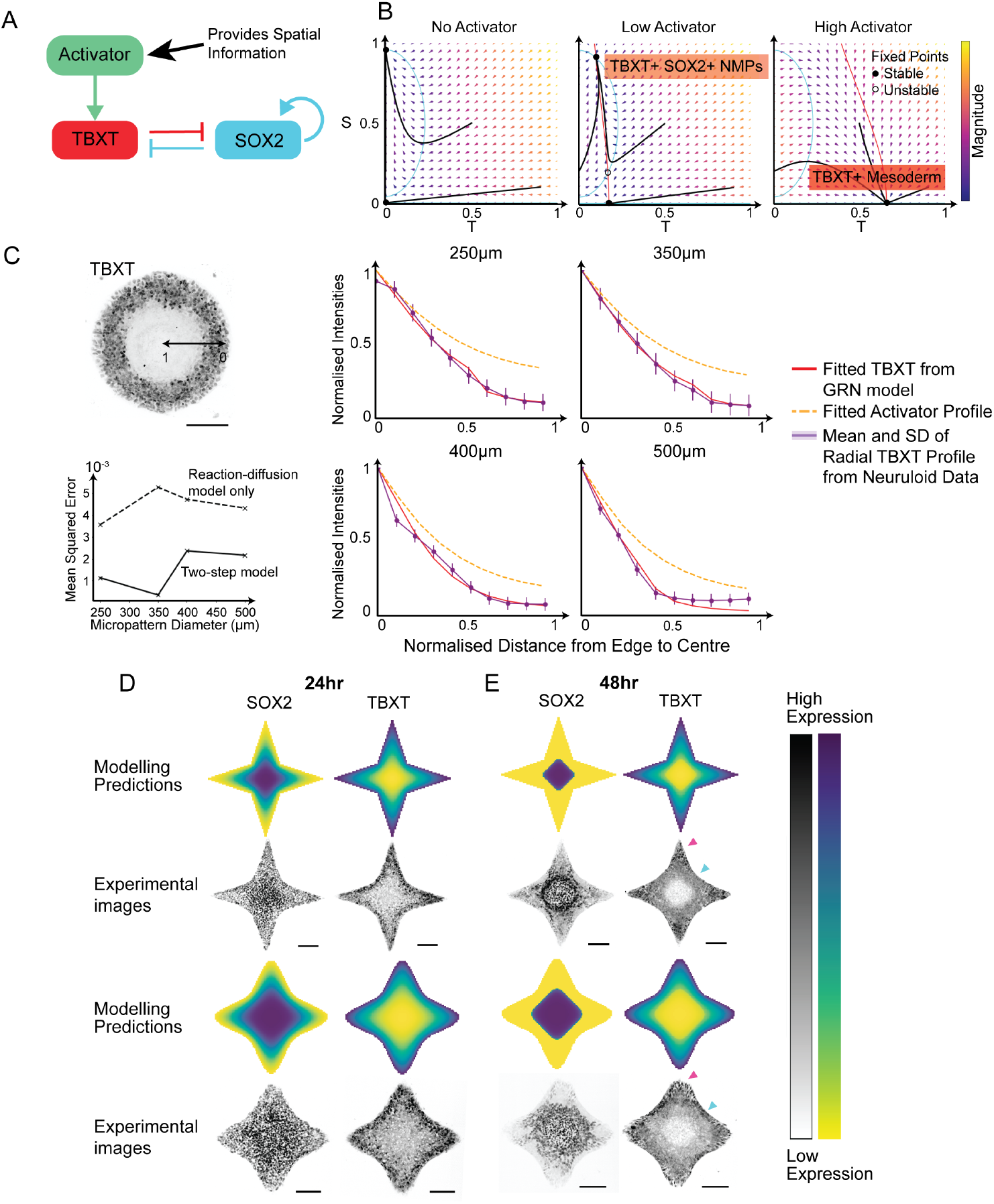
Two-step model reproduces SOX2 and TBXT profiles in varying boundary geometries. **A**. Schematic of the two-step model. The activator concentration is an input from the reaction-diffusion model (step 1) (Fig. 2), while the GRN describes the interaction between SOX2 and TBXT (step 2). **B**. Phase diagrams with gene expression trajectories for different activator levels. The system is bistable with low activator concentration, but has only one stable fixed point for high activator concentration values. **C**. Two-step model (red solid lines) fitted to TBXT intensity (purple solid lines) from neuruloid data, and corresponding activator profile (orange dashed line) required for increasing micropattern diameters. At least *n* = 3 neuruloids were used for each micropattern diameter. Lower left plot shows mean squared error between model and TBXT data for reaction-diffusion model only (dashed line) and two-step model (solid line). **D-E**. SOX2 and TBXT patterning at 24 (D) and 48 (E) hours post induction on sharp and curved domains with both convex and concave boundaries. Grayscale images show experimental data, with heatmaps at the top showing model predictions. Magenta and cyan arrows indicate convex and concave corners respectively. Scale bars: All scale bars are 100*µ*m.

In this two-step model, we assumed that chemical reactions and diffusion occurred on faster timescales than gene regulatory interactions. With this separation of time scales, we solved the reaction-diffusion model to steady-state to obtain the stable spatially-varying activator concentration. This was then input into the ODE model governing SOX2 and TBXT dynamics. We explored how the activator concentration varied the balance of SOX2 and TBXT expression levels. With varying activator concentration, the model exhibited a bifurcation behaviour, capturing bistable states at intermediate levels of activator concentrations (Fig. 4B). There existed fixed points with co-expression of SOX2 and TBXT, representing the NMP state. At higher activator concentrations, the trajectories consistently transitioned towards a mesodermal fate, characterised by high TBXT expression with no SOX2 expression (Fig. 4B).

We fitted our two-step model, using Markov chain Monte Carlo (MCMC) methods, to TBXT intensity profiles from 24-48 hour neuruloids, where spatial patterning was prominent prior to the onset of 3D morphogenesis (Fig. 4C and S5A-B and Methods). Extracting the parameters for transcription and degradation rates of SOX2 and TBXT from MCMC, our two-step model resulted in a better fit with lower mean squared error to the experimental TBXT intensity data (Fig. 4C left bottom row) compared with reaction-diffusion model which considers the activator alone, as expected. This fitting procedure also yielded an estimate of the activator concentration profile required to obtain the observed TBXT expression (Fig. 4C orange dashed lines).

To further investigate the role of boundary geometry in pattern formation, we simulated the two-step model on 2D domains with more complex boundary properties with identical parameters to those fitted from Fig. 4C. We generated “sharp star” and “curved star” domains, which include both convex and concave edges (Fig. 4D-E top panels). These geometries allowed us to compare shapes with identical surface areas, isolating the effects of vertex sharpness and curvature types (positive vs. negative). The two domains have different degrees of curvatures at the corners. The simulations show distinct spatiotemporal properties in SOX2 and TBXT expressions, where the down-regulation and up-regulation of SOX2 and TBXT intensities were more pronounced at the convex corners compared to the concave corners. Since the assumption is that signalling is primarily boundary-driven, these simulations align with our hypothesis that regions near sharper boundaries have increased sensitivity to the activator, as they received input from two adjacent boundary segments. In contrast, regions near concave boundaries would show reduced sensitivity, as cells are positioned further from the nearest boundary source.

To test the model predictions, neuruloids were generated on micropatterns with the same “sharp star” and “curved star” geometries. We examined SOX2 and TBXT patterns at 24 and 48 hours post-induction in these domains (Fig. 4D-E bottom panels). Aligned to the model predictions, we observed TBXT up-regulation within the “limbs” of the shape over time (indicated by magenta arrows in Fig. 4E). At 48 hours, this effect was more pronounced at sharp vertices, consistent with previous observations on the triangle domain (Fig. 3B and S5C-D). Comparing the TBXT intensities between sharp and wavy stars at the convex corners, and the transition from 24 to 48 hour neuruloids, the sharp stars show more spatially isolated up-regulation of TBXT in the limbs compared to the wavy star. SOX2 expression was notably down-regulated along these same regions. Consistent with our prediction, concave edges appeared to have a weaker influence on SOX2 and TBXT patterning compared to convex ones. A similar, though subtler, effect was seen in the annulus domains, where TBXT intensity was lower along the concave inner boundary than along the convex outer edge (Fig. S4C).

The model predicted spatial heterogeneity in SOX2 and TBXT expressions matching those observed in neuruloids. Given the predictive ability of our model parameterised on only a subset of the data (circular domains) to a range of different challenges with no further fitting, we were confident that the close agreement was not due to over-fitting. Both simulation and experimental results are consistent with the notion that signals from adjacent boundaries may overlap in regions with acute angles, resulting in an increase in the sensitivity to signalling due to the diffusion process. Together with the results from the previous section, these suggest that a diffusive molecular signal originating from the boundaries can account for the edge sensing mechanism in this micropatterned system, without the need for mechanical inputs^50^.

To conclude, a minimal and mutually repressive gene regulatory network, downstream of a boundary-driven activation was sufficient to capture the emergence of SOX2 and TBXT patterns arising from a variety of geometric perturbations. We note that neuruloids are a complex self-organising system, and thus there are likely other factors contributing to patterning, yet our minimal model predicted global patterns under different boundary conditions.

### Inhibition of Wnt secretion preserves initial patterning but disrupts subsequent morphological development

Our focus so far has been to develop a phenomenological model of observed cell fate patterns within neuruloids, assuming the existence of a general activator molecule. Wnt signalling has been shown to play a critical and concentration-dependent role in regulating TBXT expression^51^. Additionally, high local Wnt concentrations drive NMPs toward mesodermal fates by promoting the Wnt/FGF signalling cascade and expression of TBXT, whereas low levels of Wnt facilitate commitment to neural fates during vertebrate axial elongation^46,48,52^. Wnt and its inhibitor DKK are well-established components of an activator-inhibitor system in the canonical Wnt/*β*-catenin signalling pathway^53^, which has been previously modelled in the context of hair follicle spatial patterning^54^. Thus, we explored the possibility that the Wnt pathway may form at least part of the activator signal.

We first searched for Wnt signalling pathway components within transcriptomic data from day-3 neuruloids^23^. Significant ligand-receptor interactions involving Wnt3 and DKK ligands, inferred bioinformatically^55^, predominantly originated from mesoderm and primitive streak populations within the neuruloid system (Fig. S6A).

To visualise spatial Wnt activity, we used LEF1 as a readout of Wnt activation. As a downstream effector of the Wnt/*β*-catenin signalling pathway^53^, LEF1 serves as a proxy for local Wnt concentration. The expression of LEF1 displayed the same spatial profiles as TBXT in 24-hour neuruloids (Fig. 5A). Furthermore, the expression profiles of LEF1 (taken to represent the activator) and TBXT quantified for neuruloids of different micropattern sizes closely resembled the predictions from our two-step model (Fig. 4C and Fig. 5A). To confirm that the time scale of gradient formation was much faster than cell differentiation, we checked LEF1 expression at 8 hours post-induction (Fig. 5B-C). Under control conditions, we observed a clear LEF1 gradient without TBXT expression (Fig. 5C). The emergence of the “activator” prior to the downstream patterning is consistent with the quasi-static approximation utilised above.

**Figure 5:**
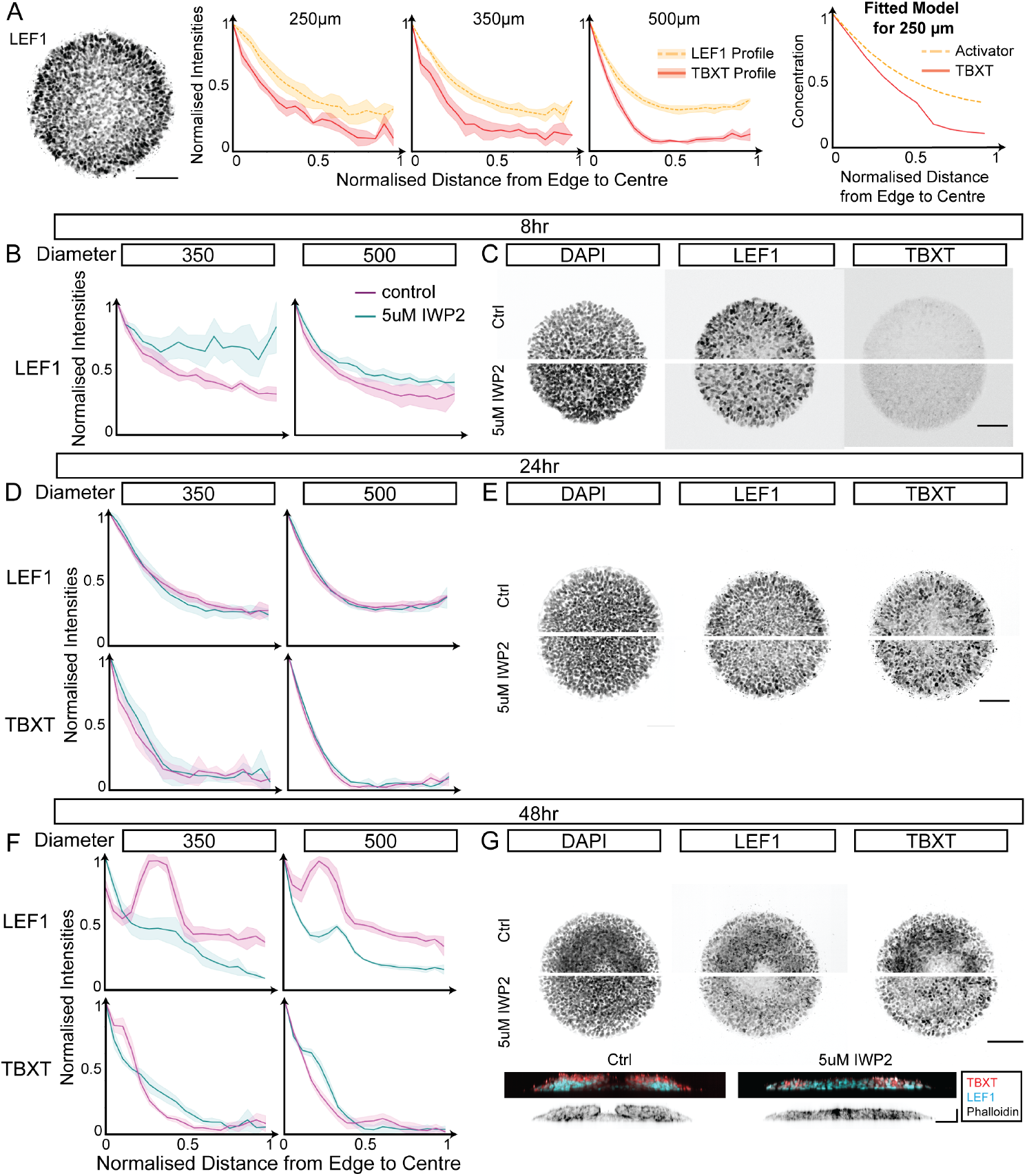
LEF1 gradient matches activator gradient from model fitting but pattern emergence was not disrupted when Wnt secretion was inhibited. **A**. LEF1 (orange dashed lines) and TBXT (red solid lines) profiles extracted from 24-hour neuruloid data for increasing micropattern diameters. At least *n* = 3 neuruloids were used for each micropattern diameters. **B, D, F**. LEF1 and TBXT intensity profiles for control (magenta) and IWP2-treated (teal) conditions for 8-hour (B), 24-hour (D) and 48 hour (F) neuruloids. At least *n* = 3 neuruloids were used for each micropattern diameters and experimental conditions. **C, E, G**. DAPI, LEF1 and TBXT signals, with control (top) and inhibited conditions (lower). **G** Lower panels show cross-sectional view of neuruloids at 48-hours. Neuruloids chosen in C, E and G had similar cell densities as detected using segmentation. Scale bars: All horizontal scale bars are 100*µ*m, vertical scale bar is 50*µ*m.

As canonical Wnt signalling is activated uniformly upon induction, we reasoned that spatial patterning might arise from endogenously secreted Wnt ligands^56^. To investigate this, we inhibited Wnt secretion using IWP2, a small molecule that blocks Porcupine-mediated Wnt ligand processing^57^. Initially, we saw a clear difference in LEF1 profiles 8 hours after induction, before the onset of TBXT (Fig. 5B-C). Surprisingly, at 24 hours post-induction, we observed no significant variations in the spatial expression of TBXT or LEF1 between control and IWP2-treated conditions (Fig. 5D-E). This suggests that pattern emergence is not immediately disrupted by interference with Wnt secretion. However, by 48 hours, TBXT and LEF1 expression profiles in IWP2-treated samples deviated markedly from controls (Fig. 5F-G), indicating a pronounced effect of Wnt inhibition on further pattern maintenance and development. Cross-sectional views at 48 hours post induction revealed striking morphological differences between the control and IWP2-treated conditions (Fig. 5G). In particular, the characteristic “doughnut” morphology in which the tissue periphery exhibit more cell layers than the centre, was present in the 48-hour control condition (Fig. 5G) and in previous reports^23^, but was absent following IWP2 treatment. This suggests that boundary-driven pattern emergence may initially rely on additional molecular cues, while Wnt signalling is crucial for proper development, shaping both morphogenesis and downstream spatial patterning.

### Reaction-diffusion description of later neuruloid development

To this point, we have focused on the role of boundaries in the emergence of TBXT patterning of neuruloids after induction with signalling molecules. Interestingly, at later stages of neuruloid development (3 days post-induction), a second symmetry-breaking event in cell fate patterning has been previously observed^23^. This occurs after morphogenetic events at 48-hour neuruloid development described in the previous section. During this phase, the initial radially uniform TBXT-positive mesodermal ring gave rise to discrete, overgrown clusters of TBXT-positive cells at the neuruloid periphery, with the number of clusters dependent on micropattern size (Figure 3B in Rito *et al*.^23^ and Fig. 6A). Since reaction-diffusion frameworks have previously successfully described such size-dependent periodic patterns, we next asked whether such a phenomenological framework can also describe this behaviour.

**Figure 6:**
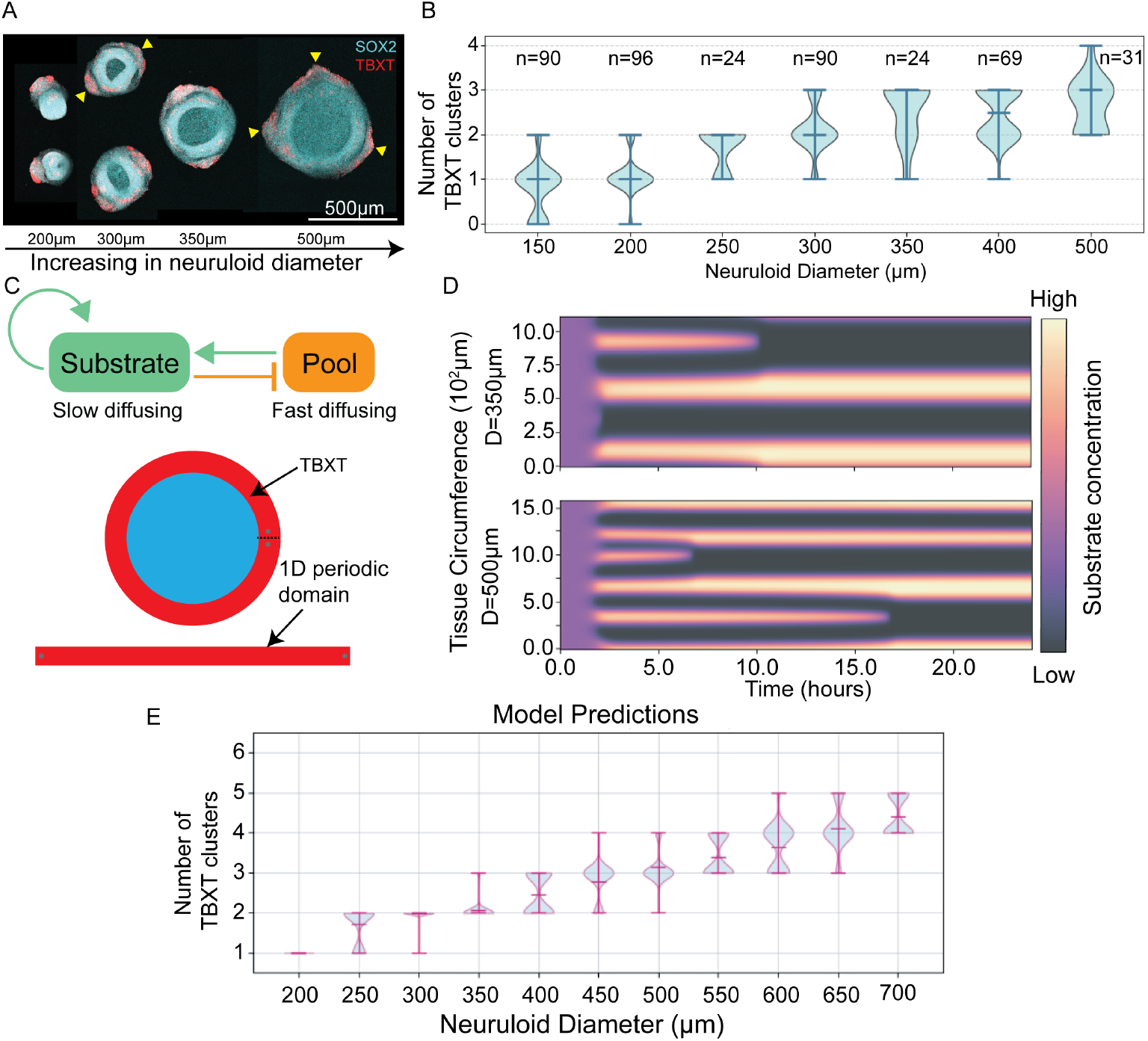
TBXT cluster formation at day 3 neuruloid development. **A**. Neuruloid images show size-dependent patterning of TBXT clusters. Yellow arrows indicate examples of the TBXT clusters. **B**. Quantification of the number of TBXT-positive cell clusters across neuruloids of increasing diameters. **C**. (Top) Schematic of substrate and pool molecule interactions in a substrate-depletion model. (Bottom) TBXT ring as a 1D periodic domain for numerical simulations. **D**. Kymograph of numerical simulations of the substrate-depletion model in increasing domain sizes, showing an increase in the number of high substrate concentration clusters with neuruloid diameter. **E**. Model predictions on the number of high TBXT expression cell clusters from *n* = 100 simulations to *T* = 24hours for increasing neuruloid diameter.

We quantified the number of these clusters across neuruloids of varying micropattern sizes by manually identifying regions that met both criteria of overgrown cellular morphology and TBXT-positive gene expression. We also measured the smallest angle between adjacent clusters (Fig. S7D-E). This analysis revealed variability in the number of clusters among neuruloids of the same size, though a general increasing trend was observed with larger micropattern diameters (Fig. 6B). Furthermore, the variability in the separation angles suggests that the formation of these clusters followed an irregular, non-uniform spatial pattern.

The transition from an initially homogeneous TBXT-positive ring to discrete clusters is reminiscent of cell polarity formation, in which localised accumulation of molecular components drives spontaneous symmetry breaking^58,59^. We hypothesised from previous results that the presence of epithelial-to-mesenchymal transition (EMT) markers in neuruloids^23^ indicate that overgrown clusters of TBXT positive cells are the result of directed cell migration guided by signalling molecules. To explore this, we utilised a phenomenological reaction-diffusion model in which the homogeneous ring of TBXT-positive cells at the neuruloid periphery is represented as a onedimensional periodic domain (Fig. 6C and see Supplementary Section S8 for details). This is taken to be the initial conditions for the simulations. The model is based on a mass-conserved substrate-depletion mechanism^60^, whereby clusters of high TBXT expression emerge through the redistribution of signalling molecules that drive TBXT-positive cells to self-organise into discrete clusters (Fig. 6C–D). This mechanism inherently generates variability in both the number and spatial positioning of clusters even in the absence of explicit stochasticity, owing to the random initial conditions. This model successfully recapitulated the experimentally observed variability in TBXT cluster number across different micropattern sizes, as well as the angular spacing between clusters (Fig. 6E; Fig. S7D–E). Together, these results demonstrate that a reaction–diffusion mechanism can also account for the second stage of symmetry breaking in TBXT patterning during neuruloid development.

## DISCUSSION

### Neuruloid pattern emergence can be quantitatively predicted from boundary-informed reaction-diffusion dynamics

We have quantitatively revealed how boundary constraints influence the spatial patterning of TBXT expression in neuruloids, an *in vitro* model recapitulating NMPs specification and their differentiation into neural and mesodermal cell types. Experimentally, by varying the size and curvatures of the micropatterned domains, we demonstrated that TBXT up-regulation is dependent on boundary geometry, exhibiting a constant length scale from the colony edge. Sharp convex curvatures led to increased TBXT expression and broader pattern widths compared with concave curvatures. Correspondingly, SOX2 down-regulation was more pronounced at convex curvatures. Within the *in vivo* progenitor zone, neuromesodermal progenitors also exhibit location-dependent fate decisions: anterior cells predominantly form neural tissue, while medial and posterior cells contribute to mesoderm^61^. Heterogeneous gene expression within confined geometries appears essential for robust axial elongation^11,12^, supporting the view that developmental patterning arises from an interplay of signalling gradients, mechanical forces, and spatial constraints rather than biochemical signals alone^62^. Although a role for confinement has been proposed, direct links between boundary conditions and fate specification remain limited. Our findings offer quantitative evidence that physical boundaries can sensitively organise gene expression patterns, influencing fate allocation in NMP populations.

The experimental TBXT patterning observed across various geometries aligns with our model in which local boundary conditions influence gene regulation, consistent with a diffusion-based mechanism. Initially, a simple diffusion-degradation model was employed to demonstrate that stable radial patterning could be fitted to the extracted TBXT spatial expression profiles (Supplementary sec. S2). However, as the domain size is increased, this model does not reproduce the formation of multiple high-intensity TBXT clusters seen in neuruloids cultured under unconstrained conditions. This limitation motivated the use of a reaction-diffusion framework instead. Moreover, the consideration of both activator and inhibitor components is supported by scRNA-seq data revealing the presence of Wnt and Dkk ligands, suggesting that their interplay underlies the observed spatial organisation. To make predictions on how this activator component regulates SOX2 and TBXT expressions which underlie NMP differentiation, to be compared directly to experimental images, we added a minimal GRN model downstream of the activator. This is, however, shown to be insufficient to fully account for pattern emergence (see final section of Discussion).

By day 3 post-induction, the initially radial TBXT pattern evolves into overgrown clusters of TBXT-positive cells at the neuruloid periphery. The number of these clusters increase with neuruloid size. This observation is consistent with classical Turing patterns and can be explained using a reaction-diffusion model with mass redistribution dynamics. Altogether, these results suggest that diffusion processes of interacting signalling molecules may underlie the emergence of both initial radial and subsequent symmetry-breaking events, which depend on domain size and boundary geometry, without requiring additional inputs.

### Mechanisms of boundary sensing in organoid systems

Micropatterned organoid models have successfully recapitulated various stages and aspects of embryonic development *in vitro*, from gastruloids that model early germ layer formation by BMP-induced signalling^21,24^ to neuruloids that use CHIR activation to study caudal cell fate patterning^23^ for embryonic trunk formation. However, these systems have each been shown to sense boundaries through distinct mechanisms. In Wnt-treated 2D gastruloids, early TBXT expression initially appeared throughout the colony before becoming edge-restricted, driven by E-Cadherin dynamics linked to radially varying cell density and *β*-catenin levels^22^. Neuruloids, despite activation of Wnt signalling using CHIR, exhibit TBXT edge restriction. This is in contrast to previous findings where activation through CHIR instead of Wnt abolished spatial patterning in micropatterned hESCs^22^. Likewise, IWP2 eliminated TBXT in 2D gastruloids^63^, but not in neuruloids, suggesting distinct sensitivities to Wnt perturbation across *in vitro* models.

While these systems exhibit distinct sensitivities to pathway perturbations, reaction-diffusion frameworks have demonstrated remarkable versatility in describing pattern emergence across diverse micropatterned systems^64,65^. This convergence suggests that reaction-diffusion mechanisms represent a fundamental principle underlying biological self-organisation^28,66^, thus providing a common theoretical framework to explain diverse patterning behaviours. Such phenomenological models not only generate testable predictions but also capture complex spatial organisation in the absence of complete molecular characterisation, as evidenced by successful applications in both developmental and skin patterning systems^38,39^.

In neuruloids, reaction-diffusion simulations across various geometries indicate that diffusion alone can account for corner-associated effects in pattern emergence. However, deviations between reaction-diffusion model predictions and experimental observations, particularly in smaller neuruloids, suggest that additional mechanisms beyond the initial patterning phase contribute to regulating the activator component and shaping the spatial distribution of TBXT. These may include morphogenetic processes associated with the onset of axial elongation. In unconstrained micropatterns, the reaction-diffusion model predicted the formation of multiple high-TBXT clusters rather than a radial pattern. While simulations yield regularly spaced clusters, experimental data reveal irregular, non-periodic spots. This discrepancy likely reflects both experimental variability and dynamic downstream regulation, consistent with the inherently non-equilibrium nature of developing neuruloids, which precludes the establishment of fully stable and predictable patterns.

In our phenomenological reaction–diffusion model, the edge effect is represented by a locally increased sensitivity to signalling at the neuruloid boundary. Lehr *et al*.^67^ implemented this effect differently, modelling a large domain in which ligand production was restricted to the micropatterned region, while diffusion and degradation occurred throughout the entire domain, including areas beyond the colony. In their framework, enhanced ligand activation was assumed at the micropattern boundary to capture edge sensing behavior. In contrast, our model assumes that the hypothesised activator, endogenously secreted Wnt, operates via short-range, intercellular signalling, and thus ligand dynamics were confined to the micropatterned domain only. However, our analyses on neuruloids with perturbed Wnt secretion indicated that Wnt may not be the sole activator of TBXT patterning. If FGF signalling also contributes to this process, the modelling approach of Lehr *et al*. may be better suited to capture the relationship between FGF and pERK signalling (discussed in final section of Discussion), particularly since FGF ligands are present in the induction medium.

### Molecular mechanisms underlying neuruloid self-organisation

Integrating the reaction-diffusion model with a minimal gene regulatory network successfully recapitulated several experimental observations, providing a rationale to investigate the underlying molecular mechanisms. The prediction of activator gradient using this model guided the search for elements of the signalling pathway underlying the patterning. In probing the molecular identity of the activator in our reaction-diffusion model, our findings suggest that Wnt signalling alone does not fully account for pattern emergence. TBXT patterning at early stages remains robust even when Wnt secretion is inhibited, though later morphogenetic processes are clearly disrupted. This result has emphasised the different roles of Wnt signalling between the initiation of patterning and spatial cell fate organisation. Since CHIR selectively activates canonical Wnt signalling, the disruption of morphogenesis following Wnt secretion inhibition suggests that additional pathways, such as FGF signalling or YAP-mediated mechanotransduction^23^, work alongside canonical Wnt activity to support both pattern emergence and morphogenesis in this system.

FGF signalling, which directly interacts with Wnt/*β*-catenin pathways, is essential for posteriorising tissues, maintaining the bipotent NMP state, and promoting mesodermal differentiation^68,69^. In *Xenopus*, FGF has been shown to regulate expression of the TBXT homolog Xbra^70^. In neuruloids, FGF/ERK signalling displays a pattern similar to TBXT^23^, and FGF receptors were recently found to localise basally on micropatterned gastruloids, akin to BMP receptors^24,71^. These findings raise the possibility that FGF ligands contribute to the molecular basis of the reaction-diffusion system underpinning TBXT patterning. Further work will better inform the specific contribution of the FGF/Erk pathway in spatial self-organisation during NMP specification.

Overall, our combined systems approach - bringing together both experimental and theoretical approaches - provides a rigorous and predictive framework for understanding how changes in boundary constraints impact tissue self-organisation.

### Limitations of the study

Experimentally, we were unable to easily visualise key components directly, such as Wnt activity. Imaging the localisation of *β*-catenin at the tissue scale was challenging with commercial antibodies. We were unable to vary concentrations of signalling molecules, Wnt and FGF, since this would alter the cell fates obtained and the axial progenitor system itself. In principle, our models were sufficient to reproduce the observed patterns and variability in the data. However, the choice of the specific reaction-diffusion model was not constrained, as our goal was to provide a phenomenological description rather than a mechanistic explanation. Consequently, the model parameters are arbitrary: they are not uniquely defined and do not necessarily correspond directly to specific experimental conditions.

## METHODS

### Experimental methods

#### Human embryonic stem cell culture

H9 human embryonic stem cells (hESCs) were cultured using StemFlex medium (Thermo Fisher Scientific; A3349401) in T25 flasks coated with 0.5 *µ*l/cm^2^ laminin-521 (Thermo Fisher Scientific; A29249). When flasks reached 75-80% confluence, cells were passaged at ratios of 1:6 and 1:8 using ReLeSR as per manufacturer’s instructions (StemCell Technologies; 05872).

All human embryonic stem cell experiments were conducted at the University of Warwick in accordance with the Guidelines for Stem Cell Research and Clinical Translation from the International Society for Stem Cell Research, as well as the UK Code of Practice for the Use of Human Stem Cell Lines. Our culture system represents a defined stage of development and adheres to section 2.2.1A of the ISSCR Guidelines.

#### Micropatterning coverslips

Micropatterns were generated following the protocol described in^72^. Briefly, IPA-treated coverslips were first activated in a UVO-cleaner for 10 min, and then incubated with PLL(20)-g[3.5]-PEG(5) for 1.5 hours. Coated coverslips were placed onto a chrome mask that had been UV-ozone activated for 2 minutes and contained the desired pattern geometries. Adherence between the coverslip and mask was aided by adding dH_2_O. The assembly was then inverted so the silver side faced upward and exposed to UV light for 8 minutes. Coverslips were subsequently immersed in 70% ethanol for 15 minutes, air-dried, and used within 4 weeks.

#### Generation of posterior neuruloids

Posterior neuruloids were produced according to^23^. In summary, micropatterned coverslips were thoroughly washed using PBS+/+ (PBS with calcium and magnesium; Thermo Fisher Scientific, 14040-091), and then incubated with rh-Laminin-521 (Thermo Fisher Scientific, A29248) diluted 1:10 in PBS+/+ for 3 hours at 37°C, or overnight at 4°C. hESCs were washed using PBS, dissociated using Accutase, and then manually adjusted to 600,000 cells/ml in StemFlex with 10*µ*M Y-27632. 1.5ml of the single-cell suspension was added to each coated coverslip in a 6-well plate well. After 3 hours, wells containing coverslips were washed once with PBS, and the medium was replaced with fresh StemFlex for overnight incubation. After 18 hours, colonies were induced using 2.5ml of induction media consisting of N2B27 media with CHIR99021 (3*µ*M, StemCell Technologies), FGF2 (5ng/mL), SB431542 (10*µ*M) and LDN193189 (0.1*µ*M). Medium was replaced again the following day. Coverslips were fixed using 4% freshly prepared PFA for 30 minutes at room temperature, then washed twice with PBS and stored at 4°C for subsequent analysis.

#### Immunostaining

Cells cultured on micropatterns and within 3D colonies were blocked and permeabilised at room temperature for one hour in a solution containing DPBS-/-, 1% Triton-X, 10% DMSO, 10% SDS, and 4% Normal Donkey Serum. After a brief rinse in DPBS-/-(1 minute), samples were incubated overnight at 4°C with primary antibodies. The following day, samples were washed three times for 5 minutes each in PBS-/-, then incubated overnight with secondary antibodies conjugated to Alexa Fluor dyes (488, 555, 594, or 647; 1:1000 dilution) and DAPI. A final 10-minute wash in DPBS-/-was performed before mounting with ProLong Glass Antifade Mountant (Invitrogen). Mounted samples were imaged using a spinning disk confocal microscope.

### Mathematical Modelling

#### Reaction-diffusion modelling on 2D domain

The true dynamics of interactions between the hypothesised activator and inhibitor component are not known. Hence, we used a general reaction-diffusion model solved in the domain Ω ∈ ℝ^2^ describing concentrations of activator and inhibitor, denoted by *A* and *I* respectively. This model was first proposed by the Kondo group^73,74^ to study patterning on the fish skin. The coupled partial differential equations (PDEs) are

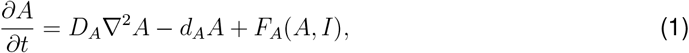

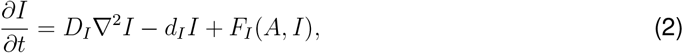

where the three terms for both equations model the diffusion, degradation and production, respectively, for both of the activator and inhibitor. *D*_*A*_ and *D*_*I*_ are diffusion coefficients; while *d*_*A*_ and *d*_*I*_ are degradation rates of activator and inhibitor molecules respectively.

We model the production functions with linear terms

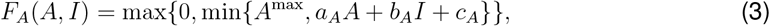

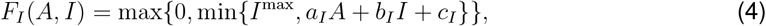

where nonlinearity is added by truncating the function. The parameters *a*_*A*_, *b*_*A*_, *c*_*A*_, *a*_*I*_, *b*_*I*_ and *c*_*I*_ are arbitrary parameters to model the interactions between activator and inhibitor. In this model, we ensure that the parameters are chosen so that they are in the diffusion driven instability regime from linear stability analysis and Turing patterns can be formed (Supplementary sec S3). A mixed boundary condition is used for this model, in the form of

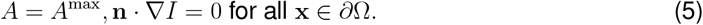

Previous analyses of mixed boundary conditions have shown this produces more symmetrical spatial patterning^43^ that is robust to fluctuations, making it an ideal framework for capturing differential responses to boundaries in a simplified, phenomenological manner. Hence, we used a Dirichlet boundary condition for the activator concentration to impose the condition that cells at the edge have an increased sensitivity to signalling molecules, as detailed in the results section^22,24,42^. The Neumann boundary condition was used for the inhibitor concentration, indicating no flow of inhibitor out of the micropattern since we hypothesised that the inhibitors are endogeneously and intracellularly produced. The equations were solved using finite element methods and sensitivity analysis is performed to determine a suitable set of parameters to model radially symmetric patterning (see Supplementary sec S3). Finite element method was implemented on Python *FEniCS* library^75^. Meshes for simulations were generated using *mshr* library according to the micropattern domains from experiments. Non-dimensionalisation and analysis of parameters is shown in Supplementary sec S3.

#### Minimal model of mutually inhibiting SOX2 and TBXT

To study the effects of activator concentration on gene regulation, we construct a minimal ordinary differential equation (ODE) model to obtain SOX2 and TBXT profile predictions. We considered the concentrations of two mutually inhibiting transcription factors SOX2 and TBXT, *S* and *T*, respectively,

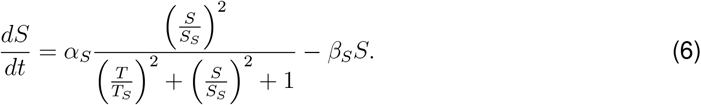

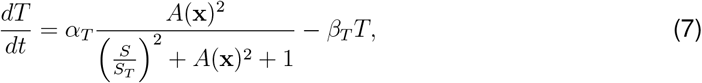

*A*(**x**) is the activator concentration which varies in spatial coordinates, **x** Ω∈ (Ω is the micropattern domain) and hypothesised to promote TBXT transcription. The model takes the stable concentration profile of activator from the reaction-diffusion model, *A*(**x**) (see previous section of Methods) as input that promotes the expression of TBXT. The activation and inhibition of gene interactions as shown in Fig. 4A are represented using Hill functions, where *S*_*S*_, *T*_*S*_ and *S*_*T*_ are scaling constants that represent the concentration when production rate is half of the maximum. *α*_*S*_ and *α*_*T*_ are maximum transcription rates; while *β*_*S*_ and *β*_*T*_ are the degradation rates of the transcription factors SOX2 and TBXT respectively. The general form for the production term of the transcription factors,

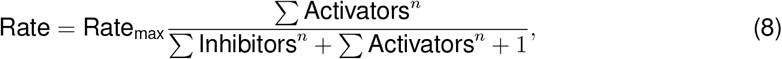

is used to model the competition between activators and inhibitors for the regulatory effects in the system.

Initial conditions for the model are *S*_0_ = 0.2 and *T*_0_ = 0 for initial SOX2 and TBXT signals respectively, since pluripotent stem cells express low levels of SOX2. The activator profile here provides spatial information to the cells, in line with the concept of positional information^30^. The nullclines of the ODE satisfy

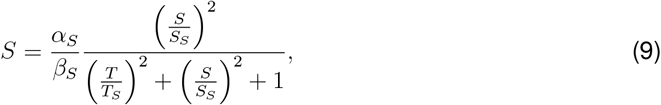

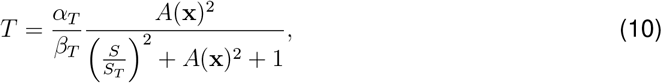

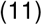

where the intersections of these equations are the fixed points of the system. The equations were solved numerically using *scipy*.*integrate*.*odeint* function on Python. Details on nondimensionalisation, stability analysis of the fixed points and the parameter space are in Supplementary sec S5.

#### Substrate-depletion model for TBXT cluster formation

To capture the TBXT-positive cell cluster formation, we model the initially homogeneous ring of TBXT expression at the neuruloid edge as a 1D periodic domain *x* ∈ [0, *L*], as illustrated in Fig. 6, where *L* is the length of the tissue circumference. We utilised a mass-conserved substrate-depletion model^60^, in which the concentration of a regulatory substrate denoted by *c*, is coupled to a finite pool of building blocks, denoted by *P*. The system is described by the following PDE

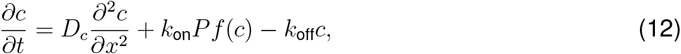

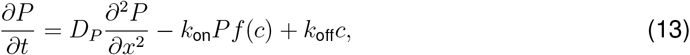

where *D*_*c*_ and *D*_*P*_ are the diffusion coefficients of the substrate and pool respectively and *D*_*c*_ ≪ *D*_*P*_. *k*_on_ represents the association rate of *P* to form *c* while *k*_off_ is the dissociation rate. Cooperative assembly is captured by the Hill 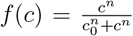, where the constant *n >* 1 controls the strength of cooperative assembly of *c*. Importantly, the total molecular content of the system, *N* is conserved at all times

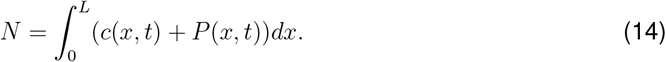

Further details on the parameters are in Supplementary sec S8.

### Image Analysis

All image analysis were done using a combination of Fiji ImageJ^76^ and custom code on Python, including using Cellpose^77,78^ for segmentation and libraries including *numpy, scikit-image*, and *matplotlib* for visualisation.

#### Analysis of fluorescence intensity

To quantify gene expression patterns for early neuruloid timepoints (day 1 post-induction), we performed analysis on fluorescence intensity based on nuclear segmentation. First, the best focal plane was selected from each 3D image stack by computing the Laplacian variance on the DAPI channel. The image slice with the largest value has the sharpest focus. All subsequent analyses were conducted on this selected plane. Nuclear segmentation was performed using a custom-trained Cellpose model^78^, fine-tuned to improve accuracy over the default nuclei model.

The segmentation masks were used to generate a binary image, from which colony contours were identified using the *scikit-image*.*measure*.*find contours* function. The colony edge is then detected from the masks of the nuclei segmentation. The centroids of each segmented nucleus was extracted from each mask and used to compute the distance from the colony edge. For each nucleus, gene expression intensity was determined by computing the mean fluorescence intensity value within the nuclear region for the corresponding channel. To account for variability in staining or imaging conditions, fluorescence intensities were normalised to DAPI signal per cell. This is done by dividing transcription factor intensities such as SOX2 and TBXT of the cell by its DAPI signal.

To generate intensity profiles as a function of distance from the colony boundary, the radial distance from the edge to the center (largest nuclei distance from edge) was discretised into *N* regions. Cell centroids located between distances *L*_*k*_ and *L*_*k*+1_, where *k* ∈ [0, *N*], were grouped and the mean intensity of each group was computed to construct a radial intensity profile. Fig S5 illustrates this analysis pipeline and the resulting quantification. For larger neuruloids where tile scans were required, contrast limited adaptive histogram equalization (CLAHE) method was done on the images before extracting the intensities to correct variation in intensities due to tiling. The same analysis was done for neuruloids cultured on geometrical micropatterns.

For later-stage and larger neuruloids, where lower resolution imaging makes accurate singlecell segmentation unreliable, we analysed fluorescence intensity from maximum intensity projections of the image stacks. To detect the neuruloid boundary in both circular and polygonal micropatterns, we applied a threshold to the DAPI channel to generate a binary image. Using this mask, we extracted the outer contour using scikit-image’s contour detection. With the boundary coordinates, we can calculate the centre of area of the contour using the formula

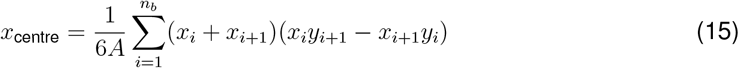

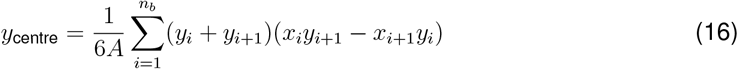

where (*x*_*i*_, *y*_*i*_) are the coordinates of the *i*^*th*^ boundary point and *n*_*b*_ is the total number of points of the contour. *A* is the area computed from the same coordinate list. For circular neuruloids, the effective radius was estimated by averaging the shortest and longest distances from the centroid to the boundary. Similarly, we can discretise the neuruloid into *N* regions based on distance to the nearest boundary. Mean fluorescence intensities were calculated for each discretised region. Similarly, the transcription factor intensity profiles can be normalised with DAPI intensity profile. A similar analysis was also applied to the simulated 2D concentration data from the reaction-diffusion model on geometrical domains, enabling direct comparison between experimental and theoretical results.

#### Parameter fitting

The models are fitted to the normalised signal intensities obtained from neuruloid images. To compare reaction-diffusion model to extracted TBXT intensities, we found parameter values that give the minimum mean absolute error (see Fig. 3A and Fig. S5C). To fit the full two-stage model to the TBXT data, we utilised Markov Chain Monte Carlo (MCMC) methods to estimate a set of parameters to fit to neuruloid data of different sizes (see Supplementary sec S6 for more details).

Since the underlying patterning mechanism remains undefined, we emphasise that both the reaction–diffusion and two-stage models are phenomenological in nature. Accordingly, our focus lies in demonstrating that the predicted trends and characteristic length scales of pattern formation are consistent with experimental observations, rather than reproducing the absolute intensity of the patterns. Moreover, the reaction–diffusion equations describe molecular concentration fields rather than TBXT gene expression intensity. Thus, the value of these models lies in their ability to capture the scaling behaviour observed in the data, rather than in achieving exact quantitative agreement.

### Single-cell RNA-seq analysis

Single-cell RNA sequencing data from day 3 neuruloids were obtained from the publicly available dataset published by Rito et al. (2025), accessible via the Gene Expression Omnibus (GEO) under accession number GSE224404^23^. Principal component analysis (PCA) and UMAP were performed using default settings, with the neighbourhood graph constructed using 15 nearest neighbours and 30 principal components. scRNA-seq data was processed using the Python toolkit SCANPY^79^, and ligand–receptor interactions were inferred with CellPhoneDB v4 using the statistical analysis method^55^.

## RESOURCE AVAILABILITY

### Lead contact

Requests for further information and resources should be directed to and will be fulfilled by the lead contact, Timothy Saunders.

### Materials availability

This study did not generate new materials.

### Data and code availability

Code for image analysis and modelling can be found publicly in the github pages here and here.

## ACKNOWLEDGMENTS

Y.T.L. is supported by EPSRC through the Mathematics of Systems II CDT at the University of Warwick (reference EP/S022244/1). J.C. and R.H. are supported by the MRC Doctoral Training Partnership in Interdisciplinary Biomedical Research at Warwick (reference MR/W007053/1). S.T., G.C., J.B., and T.E.S. were supported by an EPSRC Physics of Life grant: EP/W023075/1, Reverse Engineering Morphogenesis. J.B. and T.R. are supported by the Francis Crick Institute which receives its core funding from Cancer Research UK (CC001051), the UK Medical Research Council (CC001051), and the Wellcome Trust (CC001051); by the Engineering and Physical Sciences Research Council (UK) (EP/W023865/1) and by the Wellcome Trust (220379/D/20/Z).

## AUTHOR CONTRIBUTIONS

T.E.S, J.B. and G.C. conceived and oversaw the project. Y.T.L performed mathematical modelling and numerical simulations. Y.T.L and J.C. performed image analysis and quantification of data, with support from S.T.. J.C. generated micropatterns, cultured neuruloids, stained and performed experimental perturbations with support from R.H. and T.R.. R.H. trained Cellpose nuclei segmentation model for neuruloids. Y.T.L, J.C. and T.E.S. wrote the original draft. All authors reviewed and edited the manuscript.

## SUPPLEMENTAL INFORMATION INDEX

Document S1 with Figures S1-S8, their legends and Table 1 and 2 in a PDF.

