## Supplementary Information for "Boundary constraints can determine pattern emergence"

### S1 Analysis of robust cell fate patterning

We analysed TBXT radial patterning for neuruloid culture grown in different sizes of micropatterns (see Fig. 1C). To analyse the length scale of the TBXT patterns, we obtained raw TBXT intensities profiles from day 1-2 post-induction neuruloids, and measured the full width half maximum (fwhm) of the peaks near the edge of the colony (Fig. 1D). Fig. S3A and B show that the thickness of the TBXT regions are invariant to size of micropattern domain.

### S2 Simple diffusion-degradation model on circular domain

To model the radial patterning of TBXT intensity in a circular micropattern domain, we first tried a simple diffusion-degradation model for an activator concentration, $c$, written as

$$\frac{\partial c}{\partial t} = D\nabla^2 c - kc, \tag{1}$$

where $D$ and $k$ are the diffusion coefficient and degradation rate of the activator molecule respectively. Using polar coordinates and assuming radial symmetry for $c(r,t)$, the model can be reduced to 1D and written as

$$\frac{\partial c}{\partial t} = D\left(\frac{\partial^2 c}{\partial r^2} + \frac{1}{r}\frac{\partial c}{\partial r}\right) - kc. \tag{2}$$

Solving for the steady-state $\frac{\partial c}{\partial t} = 0$, the steady-state profile $c_s(r)$ is the solution of

$$r^2\frac{\partial^2 c_s}{\partial r^2} + r\frac{\partial c_s}{\partial r} - \frac{k}{D}r^2 c_s = 0, \tag{3}$$

which is the Bessel equation with general solution

$$c_s(r) = AI_0(\gamma r) + BK_0(\gamma r), \tag{4}$$

where $I_0$ and $K_0$ are modified Bessel functions of the first and second kind respectively with order 0. The length scale parameter, $\gamma = \sqrt{k/D}$. We consider the normalised radius so that $0 \le r \le 1$, where $r = 0$ and $r = 1$ are the centre and edge of the domain respectively. We consider the Dirichlet boundary condition $c_s(1) = 1$ and $c_s(0)$ is finite. Since $K_0$ diverges when $r \to 0$, we require $B = 0$, and hence $A = 1/I_0(\gamma)$. The solution of the concentration profile at steady state is then

$$c_s(r) = \frac{I_0(\gamma r)}{I_0(\gamma)}. \tag{5}$$

We fit this to the normalised TBXT profile to obtain the relation between length scale and micropattern radius, $R$, where $\gamma \propto R$. Fig. S4 shows fitting of eqn. 5 and numerical simulations. We obtain a steeper radially symmetric gradient on the normalised radius when increasing the domain size (i.e., increasing in $\gamma$). In this model, we do not obtain high intensity clusters when further increasing in domain size as required in neuruloid patterning experiments.

### S3 Reaction-diffusion modelling and sensitivity analysis

The reaction-diffusion system used as a phenomenological model for TBXT patterning in neuruloids as shown in the methods section is

$$\frac{\partial A}{\partial t} = D_A \nabla^2 A - d_A A + F_A(A, I), \tag{6}$$

$$\frac{\partial I}{\partial t} = D_I \nabla^2 I - d_I I + F_I(A, I), \tag{7}$$

where the three terms for both equations represent the diffusion, degradation and production functions, respectively. The production functions are

$$F_A(A, I) = \max\{0, \min\{A^{\max}, a_A A + b_A I + c_A\}\}, \tag{8}$$

$$F_I(A, I) = \max\{0, \min\{I^{\max}, a_I A + b_I I + c_I\}\}, \tag{9}$$

where nonlinearity is added by truncating the function. A mixed boundary condition is used for this model, in the form of

$$A = A^{\text{bound}}, \mathbf{n} \cdot \nabla I = 0 \text{ for all } \mathbf{x} \in \partial\Omega. \tag{10}$$

We non-dimensionalised the system for a more systematic parameter search. The model for the dimensionless concentrations $\tilde{A}, \tilde{I}$, in terms of dimensionless space, $\tilde{x}$ and time $\tau$

$$\frac{\partial \tilde{A}}{\partial \tau} = \frac{\partial^2 \tilde{A}}{\partial \tilde{x}^2} + \gamma \left( \tilde{F}_{\tilde{A}}(\tilde{A}, \tilde{I}) - \tilde{A} \right), \tag{11}$$

$$\frac{\partial \tilde{I}}{\partial \tau} = \delta \frac{\partial^2 \tilde{I}}{\partial \tilde{x}^2} + \gamma \left( \tilde{F}_{\tilde{I}}(\tilde{A}, \tilde{I}) - \kappa \tilde{I} \right), \tag{12}$$

where

$$\tilde{F}_{\tilde{A}}(\tilde{A}, \tilde{I}) = \frac{1}{d_A} \max\left\{0, \min\left\{\frac{A^{\max}}{A^{\text{bound}}}, a_A \tilde{A} + b_A \tilde{I} + \frac{c_A}{A^{\text{bound}}}\right\}\right\}, \tag{13}$$

$$\tilde{F}_{\tilde{I}}(\tilde{A}, \tilde{I}) = \frac{1}{d_A} \max\left\{0, \min\left\{\frac{I^{\max}}{A^{\text{bound}}}, a_I \tilde{A} + b_I \tilde{I} + \frac{c_I}{A^{\text{bound}}}\right\}\right\}. \tag{14}$$

(15)

The boundary conditions are now

$$\tilde{A} = 1, \mathbf{n} \cdot \nabla \tilde{I} = 0 \text{ for all } \tilde{\mathbf{x}} \in \partial\Omega, \tag{16}$$

as the variables are scaled by the activator boundary value, $A^{\text{bound}}$. The dimensionless parameters are

$$\delta = \frac{D_I}{D_A}, \gamma = \frac{L^2 d_A}{D_A}, \kappa = \frac{d_I}{d_A}. \tag{17}$$

We can set $d_A = 1$, and re-scale the system (eqn 11-14) so that the production terms can now be re-written as

$$\tilde{F}_{\tilde{A}}(\tilde{A}, \tilde{I}) = \max\left\{0, \min\left\{\tilde{A}^{\max}, a_A^* \tilde{A} + b_A^* \tilde{I} + c_A^*\right\}\right\}, \tag{18}$$

$$\tilde{F}_{\tilde{I}}(\tilde{A}, \tilde{I}) = \max\left\{0, \min\left\{\tilde{I}^{\max}, a_I^* \tilde{A} + b_I^* \tilde{I} + c_I^*\right\}\right\}. \tag{19}$$

The fixed-points for activator and inhibitor concentrations of the dimensionless system, denoted by $A_{eq}, I_{eq}$ respectively, are values that satisfy

$$a_A^* A_{eq} + b_A^* I_{eq} + c_A^* - A_{eq} = 0, \tag{20}$$

$$a_I^* A_{eq} + b_I^* I_{eq} + c_I^* - \kappa I_{eq} = 0. \tag{21}$$

For the rest of the section, we will analyse the dimensionless model and drop the $^*$ in parameters for clarity. Solving the simultaneous equation, the analytical functions for the fixed points are

$$A_{eq} = \frac{-b_A c_I - c_A(\kappa - b_I)}{(a_A - 1)(\kappa - b_I) + b_A a_I}, \tag{22}$$

$$I_{eq} = \frac{c_I(a_A - 1) - a_I c_A}{(a_A - 1)(\kappa - b_I) + b_A a_I}. \tag{23}$$

To ensure Turing patterns, we use parameters that satisfy Turing diffusion driven instability, where fixed points are stable in the absence of diffusion. The linearised system is

$$\frac{d}{dt} U = AU, \tag{24}$$

where $A = \begin{pmatrix} a_A - 1 & b_A \\ a_I & b_I - \kappa \end{pmatrix}$, considering diffusion term and the Fourier modes of

$$\partial_t U = D \Delta U + AU \tag{25}$$

where $D = \begin{pmatrix} 1 & 0 \\ 0 & \delta \end{pmatrix}$. The Turing conditions for parameters in this model are

$$a_A - 1 + b_I - \kappa < 0, \tag{26}$$

$$-(a_A - 1)(\kappa - b_I) - b_A a_I > 0, \tag{27}$$

$$\delta(a_A - 1) + (b_I - \kappa) > 0, \tag{28}$$

$$(\delta(a_A - 1) + (b_I - \kappa))^2 > 4\delta((a_A - 1)(b_I - \kappa) - b_A a_I) \tag{29}$$

Satisfying these conditions, the default parameters adapted from[1], used are

| Parameters | Values |
|---|---|
| $\delta$ | 20 |
| $a_A$ | 8/3 |
| $b_A$ | -8/3 |
| $c_A$ | 1/18 |
| $a_I$ | 10/3 |
| $b_I$ | 0 |
| $c_I$ | -1/6 |
| $\kappa$ | 2 |
| $\tilde{A}^{\text{max}}$ | 2 |
| $\tilde{I}^{\text{max}}$ | 5 |

Table 1: Default parameters for dimensionless reaction-diffusion model.

The fixed points are $A_{eq} = 0.1$ and $I_{eq} = 0.083$. Fig. S3E shows sensitivity analysis plots, varying parameters $A_{eq}$, $I_{eq}$, $\gamma$ and $\delta$. The parameter $A_{eq}$ is changed by controlling the values of $c_A$ and $c_I$, while still satisfying the Turing conditions. Through numerical simulations, we found that these are the main parameters that are changing the pattern phenotypes. Effectively, $A_{eq}$ is the ratio of stable state value to the fixed boundary concentration; $\gamma$ is directly proportional to the area of the domain; $\delta$ is the ratio of diffusivity of activator to inhibitor molecules. The dimensionless model is solved using finite element methods using solver *FEniCS* on Python on a unit radius mesh.

We found that the parameter regimes where stable activator concentration $A_{eq} < 1$ and $\delta \sim 10$ resulted in robust radially symmetric patterning at small characteristic length scale, $\gamma$, with a positive concentration gradient from the centre to the boundary of the domain. When $\delta$ is large ($\sim 10^2$), high concentration clusters around the boundary are formed instead, with increasing number as domain size (or $\gamma$) increases. Based on these simulations, we fixed the parameter set listed in Table 1 for fitting to TBXT intensities, while varying $\gamma$ to fit to the TBXT data from micropatterns of varying sizes. As shown in Fig. 2B (left), increasing $\gamma$ beyond the regime that produces radial symmetry leads to the emergence of spot-like clusters of high activator concentration at the centre of the domain. The number of these clusters increases with larger $\gamma$, consistent with classical Turing instability theory. This prediction is supported by experimental data. When neuruloids were cultured in unconstrained domains, we observed clusters of high TBXT-expressing cells forming in the interior (see Fig. 2D. We quantified the cluster sizes by thresholding the TBXT intensity value to generate a binary image. We then detected the clusters using *scikit-image* and measure their axis lengths. Fig. 2D show the distribution of the axis sizes of the clusters, which are similar in range to the TBXT ring widths in the radially symmetric patterning, further supporting the existence of a characteristic length scale in TBXT patterning.

### S4 Comparing reaction-diffusion model to radially symmetric TBXT intensities circular neuruloids

From the analysis in Section S3, we established that $\gamma \propto L^2$, where $L$ represents the characteristic length scale of the system, such as the diameter of the micropattern. To compare the reaction-diffusion model with experimental TBXT intensity profiles, we varied $\gamma$ and computed the mean squared error (MSE) between the simulated activator concentration and the experimental TBXT intensity. The $\gamma$ value corresponding to the minimum MSE was taken as the best fit for each neuruloid size. This analysis revealed a clear trend between the best-fitted $\gamma$ and the squared of neuruloid diameter (see Fig. S5C), consistent with the theoretical scaling. TBXT intensity profiles were obtained through nuclear segmentation on DAPI images and quantification of corresponding fluorescence intensities from other channels, as described in the Methods section.

### S5 A minimal gene regulatory network model

We model the transitions of gene expression levels of the biprogenitor, neuromesodermal cells as a minimal ordinary differential equation (ODE) system. We model the mutually inhibiting transcription factors SOX2 and TBXT, $S$ and $T$ by writing

$$\frac{dS}{dt} = \alpha_S \frac{\left(\frac{S}{S_S}\right)^2}{\left(\frac{T}{T_S}\right)^2 + \left(\frac{S}{S_S}\right)^2 + 1} - \beta_S S, \tag{30}$$

$$\frac{dT}{dt} = \alpha_T \frac{A(\mathbf{x})^2}{\left(\frac{S}{S_T}\right)^2 + A(\mathbf{x})^2 + 1} - \beta_T T. \tag{31}$$

$$\tag{32}$$

We non-dimensionalised the equations to obtain the rescaled model

$$\frac{d\tilde{S}}{d\tau} = \frac{\left(\frac{\tilde{S}}{S_0}\right)^2}{\left(\frac{\tilde{T}}{T_0}\right)^2 + \left(\frac{\tilde{S}}{S_0}\right)^2 + 1} - \tilde{S}, \tag{33}$$

$$\frac{d\tilde{T}}{d\tau} = \alpha_0 \frac{A^2}{\left(\frac{\tilde{S}}{S_1}\right)^2 + A^2 + 1} - \beta_0 \tilde{T}, \tag{34}$$

$$\tag{35}$$

for dimensionless SOX2 and TBXT intensities, $\tilde{S}$ and $\tilde{T}$ respectively across dimensionless time $\tau$, where the dimensionless parameters are

$$\alpha_0 = \frac{\alpha_T}{\alpha_S}, \beta_0 = \frac{\beta_T}{\beta_S}, S_0 = \frac{S_S \beta_S}{\alpha_S}, S_1 = \frac{S_T \beta_S}{\alpha_S}, T_0 = \frac{T_S \beta_S}{\alpha_S}. \tag{36}$$

To analyse the stability of the equations, the nullclines (dropping tilde for clarity) for the stable state of the equations are:

$$S = \frac{\left(\frac{S}{S_0}\right)^2}{\left(\frac{T}{T_0}\right)^2 + \left(\frac{S}{S_0}\right)^2 + 1}, \tag{37}$$

$$T = \frac{\alpha_0}{\beta_0} \frac{A^2}{\left(\frac{S}{S_1}\right)^2 + A^2 + 1}, \tag{38}$$

The intersections of the curves that satisfy eqn 37 and 38 are points $(S^*, T^*)$. $S = 0$ solves eqn 37. For non-zero $S$, we can rearrange eqn 37 to obtain explicit function

$$T = T_0 \sqrt{\frac{S}{S_1^2}(1 - S) - 1}. \tag{39}$$

We bound the $S$ and $T$ values within $[0, 1]$ to allow comparison with the normalised intensity signals from neuruloid images. The nullclines for a set of parameter in Table 2 is illustrated in Fig. S7A. This figure shows that for low value of $A$ the system may converge to co-expression of TBXT and SOX2 or converge to low TBXT and no SOX2 expression, indicating neuromesodermal fates as a stable state. For higher values of A, the system will converge to high TBXT and no SOX2 expressions, indicating differentiation to mesodermal fate. The parameters for the figures are

| Parameters | Values |
|---|---|
| $\alpha_0$ | 2 |
| $\beta_0$ | 1 |
| $S_0$ | 1 |
| $S_1$ | 0.2 |
| $T_0$ | 0.1 |

Table 2: Default parameters for ODE model, to generate bifurcation plot for S7.

### S6 Fitting ODE parameters to TBXT data

To fit the two-step model to spatial TBXT intensity profiles, we used Markov Chain Monte Carlo (MCMC) method on the parameter set, $\theta = (\gamma, S_0, T_0, S_1)$. The parameters $\alpha_0$ and $\beta_0$, which control only the timescale of the gene expression trajectories and not the shape of the steady-state, were fixed to default values. For a range of discrete $\gamma$ values, we first simulated the reaction-diffusion model to obtain corresponding activator gradients as input to the gene regulatory network (GRN) model. We ensure that the model is in the parameter regime where fixed points bifurcate when activator concentration, $A$ varies between $[0, 1]$; and varied $S_0$, $T_0$, and $S_1$ to identify a parameter values that qualitatively matched the TBXT expression profiles by eye. Based on this, we constructed prior distributions as truncated normal distributions centered around these manually selected values. Fig. S7B shows the resulting prior and posterior distributions after MCMC fitting. The inset shows the MCMC trace plots, indicating convergence of the sampled parameters. MCMC was done using *PyMC* library on Python, which uses No-U-turn sampler (NUTS)[2]. The likelihood function

$$\mathcal{P}(y_{\text{data}}|\theta) = -0.5 \sum_{i=1}^{N} \left( \frac{y^i_{\text{data}} - y^i_{\text{simulated}}}{\sigma} \right)^2, \tag{40}$$

was used, where $N$ is the number of discretised regions from edge to centre of neuruloid, and $y_{\text{data}}$ and $y_{\text{simulated}}$ are the extracted TBXT intensity profile and simulated profile from the parameter set $\theta$. $\sigma$ is the mean standard deviation of the TBXT intensity profiles in space, which was obtained from multiple neuruloid data.

### S7 Analysis on scRNA-seq data

We analysed the scRNA-seq dataset of neuruloid at day 3 post-induction provided in[3]. The analysis showed activity of ligand-receptor complex interactions between Wnt and Dkk at presomitic mesoderm fates[4] (Fig. S6).

### S8 Modelling of TBXT cluster formation in late neuruloid development

At later stages of the neuruloid development, the neuruloids undergo a second symmetry-breaking event in cell fate patterning, where clusters of overgrown TBXT-expressing cells form at the periphery of the neuruloid. Fig. S1A show an example reproduced from Rito *et al.*[3]. Fig. S1B left shows the quantification of the TBXT cluster numbers across neuruloids grown on circular micropatterns of increasing diameter sizes. Due to variability in image quality, the clusters were manually identified based on the combined criteria of overgrown cell morphology and TBXT-positive gene expression. For higher resolution images, we were also able to quantify the smallest angular separation between adjacent clusters (Fig. S1B right) as illustrated schematically in Fig. S1A by yellow markers. The homogeneous ring of mesodermal cells appear to reorganise and polarise the tissue. Based on these observations and quantification, we simulated a minimal phenomenological model that recapitulated the emergence of TBXT cluster, utilising a mass-conserved substrate-depletion model. The concentration of a regulatory substrate, denoted by $c(x,t)$, is coupled to a finite pool of building blocks, denoted by $P(x,t)$ (Fig. S1C). We describe the system by the following coupled PDE

$$\frac{\partial c}{\partial t} = D_c \frac{\partial^2 c}{\partial x^2} + k_{\text{on}} P f(c) - k_{\text{off}} c, \tag{41}$$

$$\frac{\partial P}{\partial t} = D_P \frac{\partial^2 P}{\partial x^2} - k_{\text{on}} P f(c) + k_{\text{off}} c, \tag{42}$$

where $D_c$ and $D_P$ are the diffusion coefficients of the substrate and pool respectively ($D_c \ll D_P$). The term $k_{\text{on}}$ represents the association rate of $P$ to form $c$, while $k_{\text{off}}$ is the dissociation rate. Cooperative assembly is captured by the Hill function $f(c) = \frac{c^n}{c_0^n + c^n}$, where the constant $n > 1$ controls the strength of cooperative assembly of $c$[5]. Importantly, the total molecular content of the system, $N$ is conserved at all times

$$N = \int_0^L (c(x,t) + P(x,t))dx. \tag{43}$$

In this framework, pattern formation arise in this system from mass redistribution dynamics, a mechanism analogous to models widely used to describe cell polarity[6].

To investigate the spatial patterns formed by this model, we performed parameter search by numerically solving the coupled equations across a wide range of parameter values. The initial conditions used were homogeneous concentrations with random perturbations for substrate concentration $c$, and zero for pool concentration $P$ (Fig. S1D). For each parameter set, we analysed the resulting steady state profiles to identify regimes that produced clustered patterns, where the number of clusters scales with the length of the periodic domain $L$. This analysis revealed parameter regimes where the model reproduced domain size–dependent clustering, consistent with the experimental quantification (Fig. S2A-C). The number of stable clusters obtained were quantified. The predicted scaling of cluster numbers with domain size is summarised in Fig. S2D, showing good agreement with the experimental quantifications in Fig. S1B. The parameters used for simulations in Fig. S2B are shown in Table 3. In these simulations, the total concentration was kept constant for different domain sizes, where $N/L = 10$.

The results indicate that the second-stage symmetry-breaking event in neuruloids can be accounted for by a reaction–diffusion patterning mechanism, in which patterns emerge through the redistribution of signaling molecules within a mass-conserved system. Importantly, the number of symmetry-breaking nodes scales with micropattern size, revealing a characteristic length

| Parameter | Value | Units |
|---|---|---|
| $D_c$ | 0.543 | $10^4\mu\text{m}^2\text{hr}^{-1}$ |
| $D_P$ | 10.86 | $10^4\mu\text{m}^2\text{hr}^{-1}$ |
| $k_{\text{on}}$ | 250 | $\text{hr}^{-1}$ |
| $k_{\text{off}}$ | 12.5 | $\text{hr}^{-1}$ |
| $c_0$ | 20 | $10^{-2}\mu\text{m}^{-1}$ |

Table 3: Default parameters and units for the substrate-depletion model of TBXT cluster formation.

scale for cluster formation and supporting the hypothesis that molecular diffusion drives this patterning process. These observations motivate the application of a reaction–diffusion framework to model the initial radial patterning of TBXT in neuruloids.
