## Supplementary Figures for "Boundary constraints can determine pattern emergence"

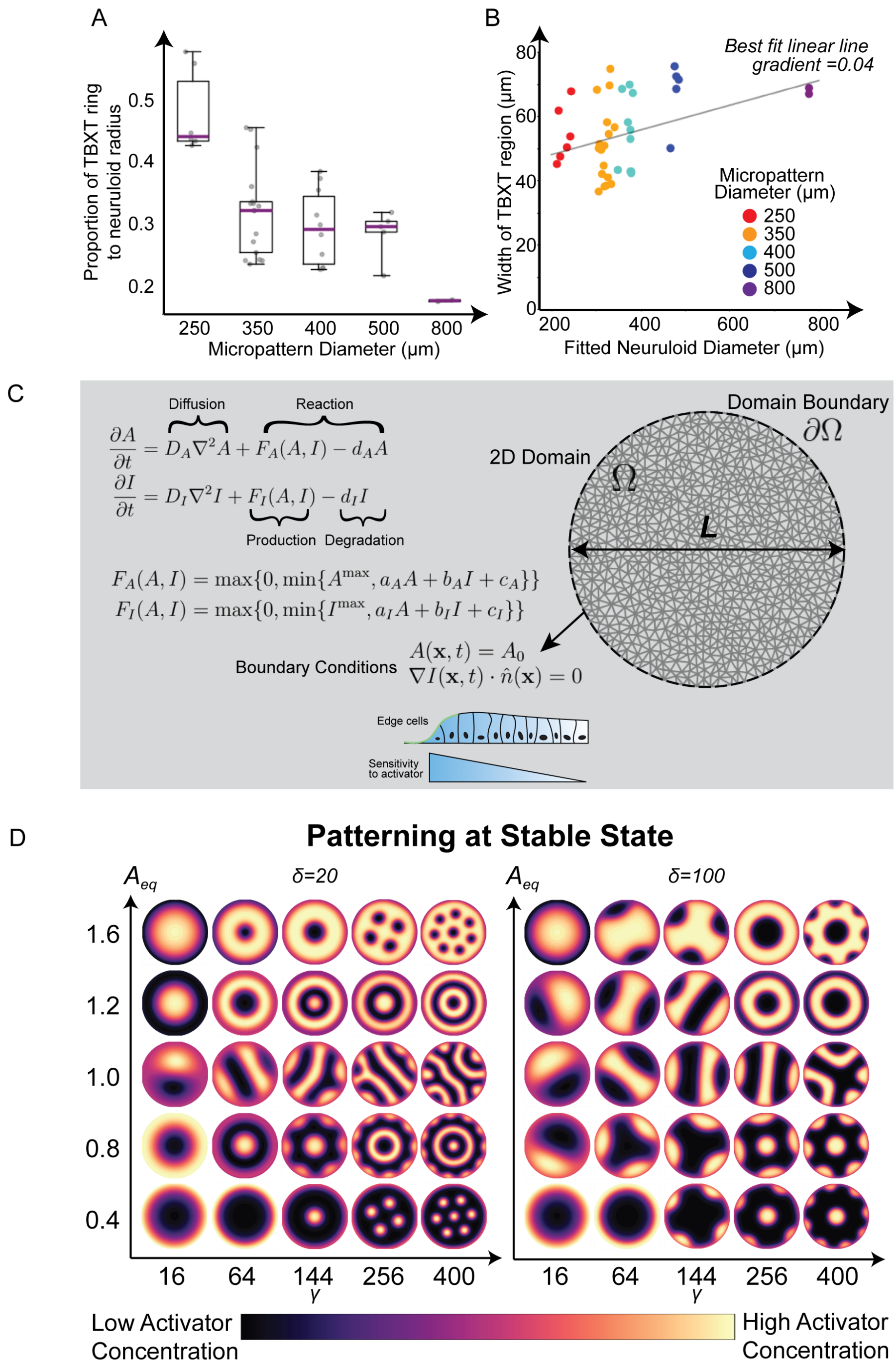


$$\frac{\partial A}{\partial t} = D_A \nabla^2 A + F_A(A,I) - d_A A$$

$$\frac{\partial I}{\partial t} = D_I \nabla^2 I + F_I(A,I) - d_I I$$

$$F_A(A,I) = \max\{0, \min\{A^{\max}, a_A A + b_A I + c_A\}\}$$

$$F_I(A,I) = \max\{0, \min\{I^{\max}, a_I A + b_I I + c_I\}\}$$

$$A(\mathbf{x},t) = A_0$$

$$\nabla I(\mathbf{x},t) \cdot \hat{n}(\mathbf{x}) = 0$$

Figure S1: ***Reaction-diffusion model predicts TBXT radial pattern.*** **A.** The ratio of the TBXT ring width to micropattern diameter decreases with increasing diameter. The boxplot displays the interquartile range (IQR), with a line indicating the median. The whiskers extend from the box to the most extreme data points lying within 1.5x the (IQR). **B.** Linear line fitted to width of TBXT region show that there is a small and insignificant trend, suggesting that the patterning is invariant to micropattern domain size. **C.** Reaction-diffusion model equations, involving diffusion, production and degradation terms. A fixed boundary condition for activator concentration is used as a minimal condition to model mechanosensation of cells to the boundary. **D.** Sensitivity analysis of reaction-diffusion model. Heatmap shows pattern formation of activator concentration at stable states for varying dimensionless parameters. $x$ and $y$-axes are dimensionless parameters $\gamma$ and $A_{eq}$ respectively. Left and right panels are simulations with different $\delta$ values (see Section S3 for parameter details).

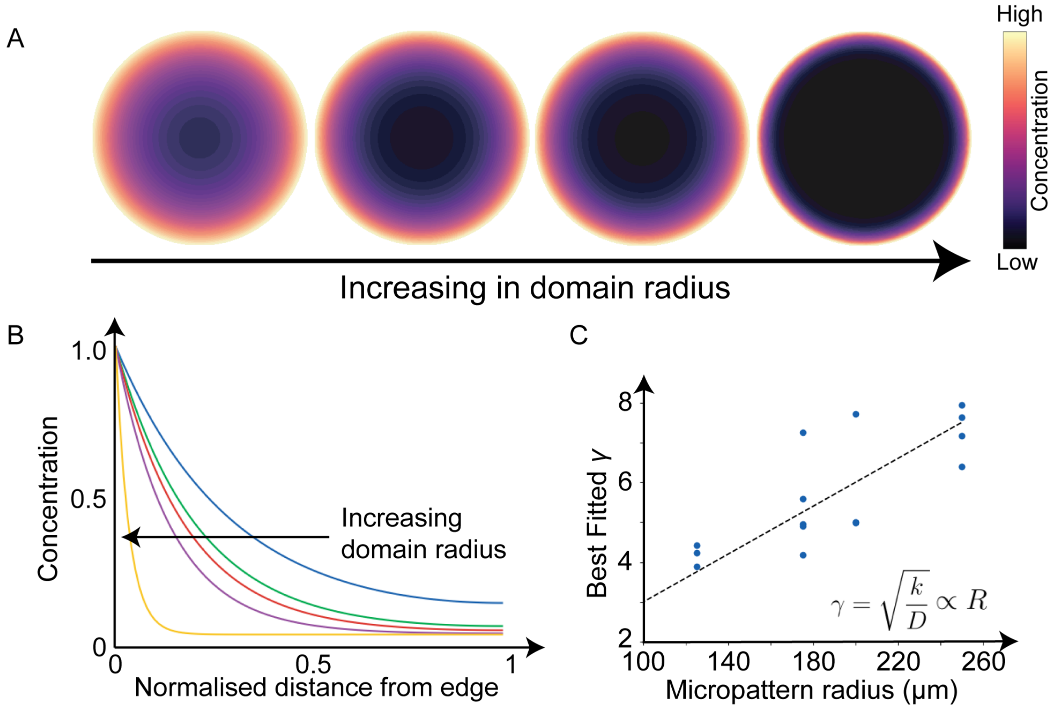


Figure S2: ***Simple diffusion-degradation model for radial patterning.*** **A.** A simple diffusion-degradation model with constant boundary conditions reproduces radial patterning for domains of various sizes (concentrations plotted on normalised domain sizes). **B.** The stable concentration, $c_s(r) = I_0(\gamma r)/I_0(r)$ show increasing steepness of gradient profile plotted on a normalised distance. **C.** Best fitted $\gamma$ obtained for TBXT intensities from neuruloids of various micropattern radii (scatter plots) and best fitted linear line (dashed line). (Section S2 for details)

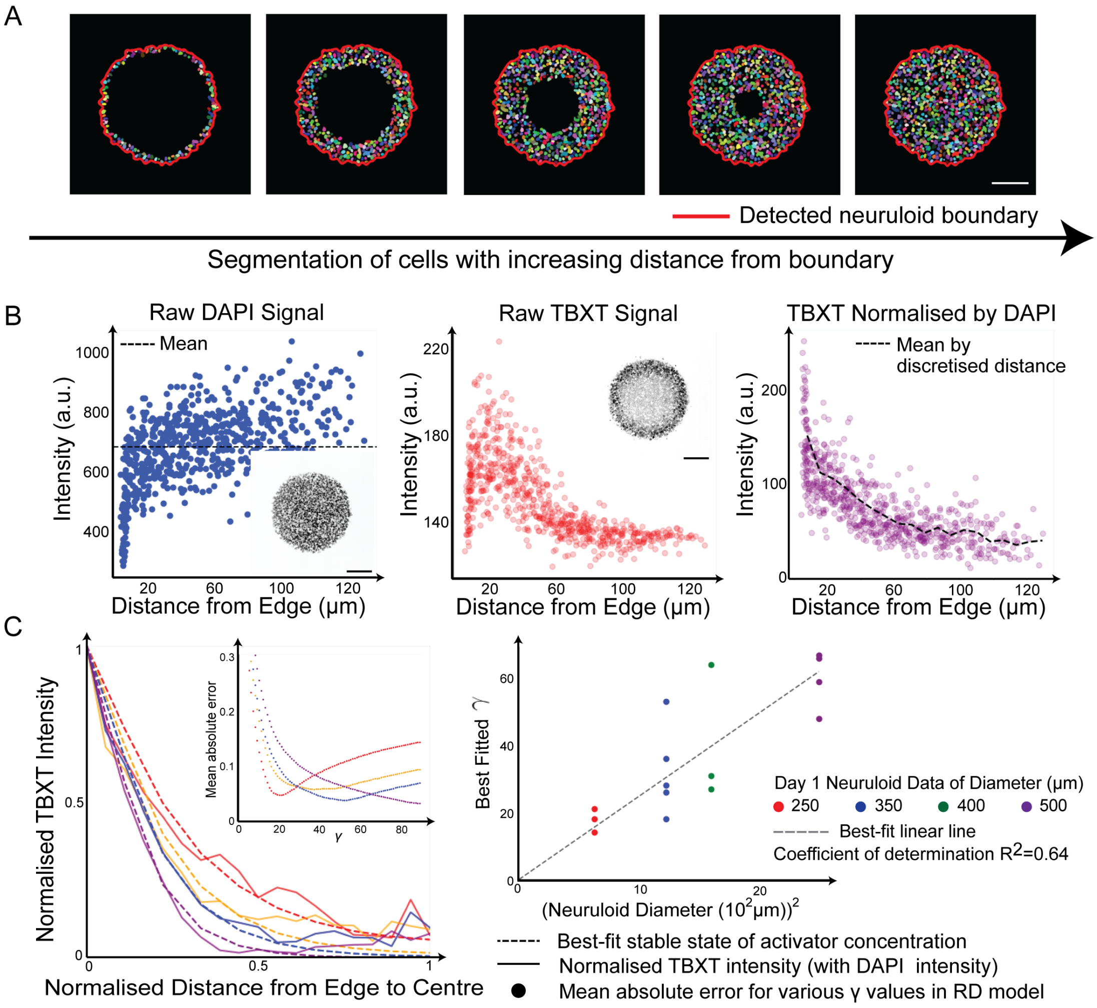


Figure S3: ***Analysis of TBXT patterns in unconstrained and circular micropatterns.*** **A.** Segmentation of circular neuruloid was done on the most focussed slice from 3D z-stack for each neuruloid image. Red curve shows the detected neuruloid boundary. Left to right panels show discretisation of distance from edge to centre of neuruloid. **B.** Raw mean intensities of DAPI (blue) and TBXT (red) for each segmented region (cell nuclei) plotted against distance from nearest neuruloid edge. Transcription factor intensities (*e.g.*, TBXT) were normalised using DAPI (purple). To generate a curve of intensity versus distance from the neuruloid edge (black dashed line), the distance from the edge to the center was discretised into intervals. Cell ROIs within each interval were grouped to obtain the mean intensity. **C.** Normalised TBXT expression (solid lines) and the best fit activator concentration gradient (dashed lines) determined by mean squared error. The best fitted $\gamma$ obtained for multiple neuruloid sizes (scatter plots) was used to fit a linear line against micropattern diameter (Section S3) to obtain the scale factor to make predictions. All scale bars are 100 $\mu$m.

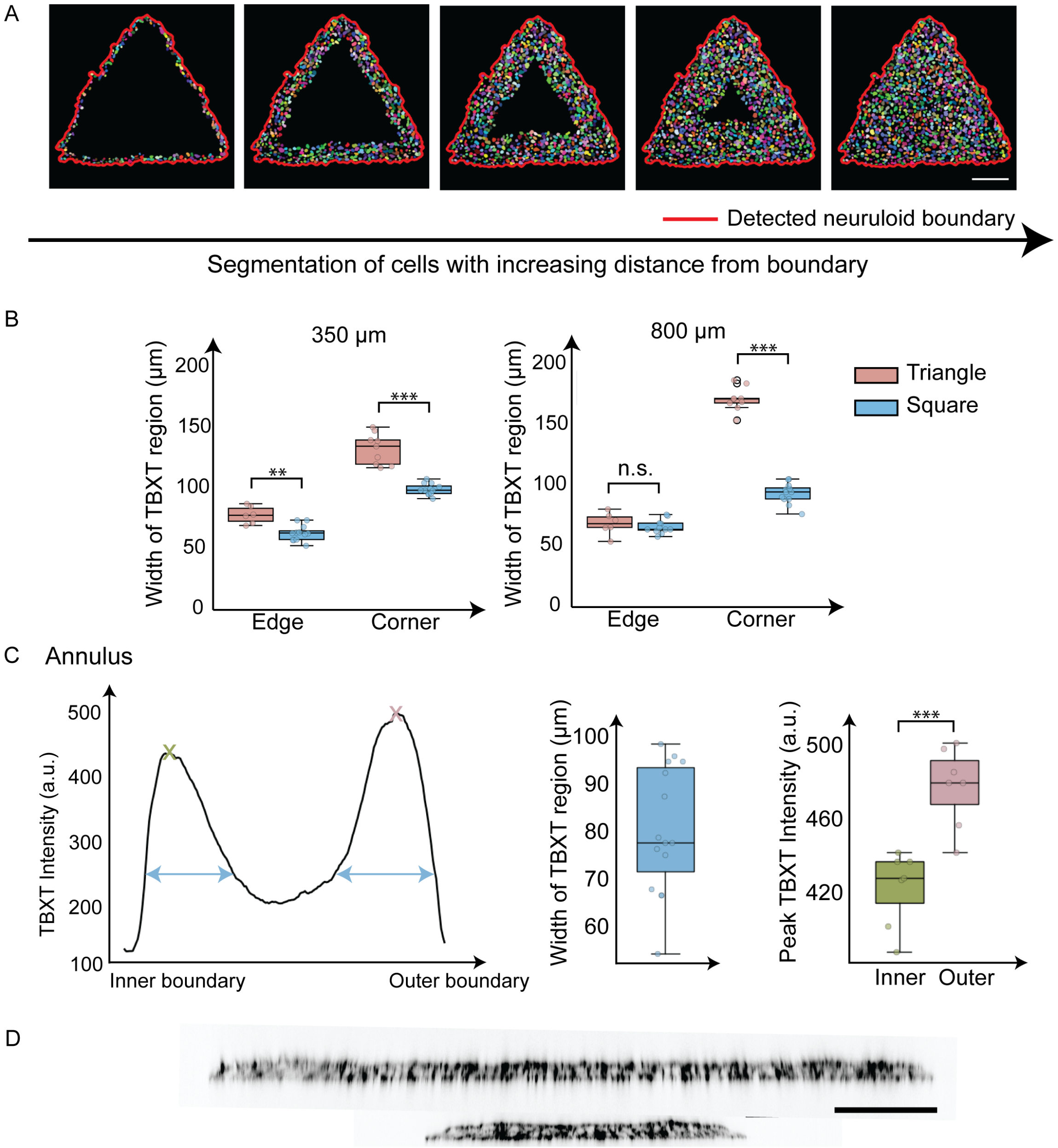


Figure S4: ***Normalised TBXT intensities from neuruloid data show good fit to reaction-diffusion model.*** **A.** Segmentation and analysis of geometrical neuruloids (triangle as example) was done with similar method to circular neuruloids. Red curve shows the detected neuruloid boundary. Left to right panels show examples of discretising the distance from edge to centre of neuruloid. **B.** Widths of TBXT pattern for triangle and square micropatterns for both edge and corners for day 2 neuruloids. There are statistically significant differences in the TBXT pattern widths at both the edges (t-test, t=4.359, p-value=0.00172) and corners (t-test, t=7.479, p-value$< 10^{-3}$) for micropattern area of 350$\mu$m diameter circle. There is no statistically significant differences at the edges (t-test, t=0.619, p-value=0.555), but statistically significant differences at the corners (t-test, t=19.034, p-value$< 10^{-3}$) for micropattern area of 800$\mu$m diameter circle. There are also statistically significant differences between edge and corner of the same shape (t-tests: 350$\mu$m triangle, t=10.55, p-value$< 10^{-3}$; 350$\mu$m square, t=15.75, p-value$< 10^{-3}$; 800$\mu$m triangle, t=20.54, p-value$< 10^{-3}$; 800$\mu$m square, t=9.245, p-value$< 10^{-3}$). **C.** Quantifying the widths of TBXT pattern (blue box plot) and the peak intensities for inner (green) and outer (pink) edge of the annulus. The difference in peak intensities was statistically significant (t-test, t=-4.905, p-value$< 10^{-3}$). All scale bars are 100 $\mu$m.

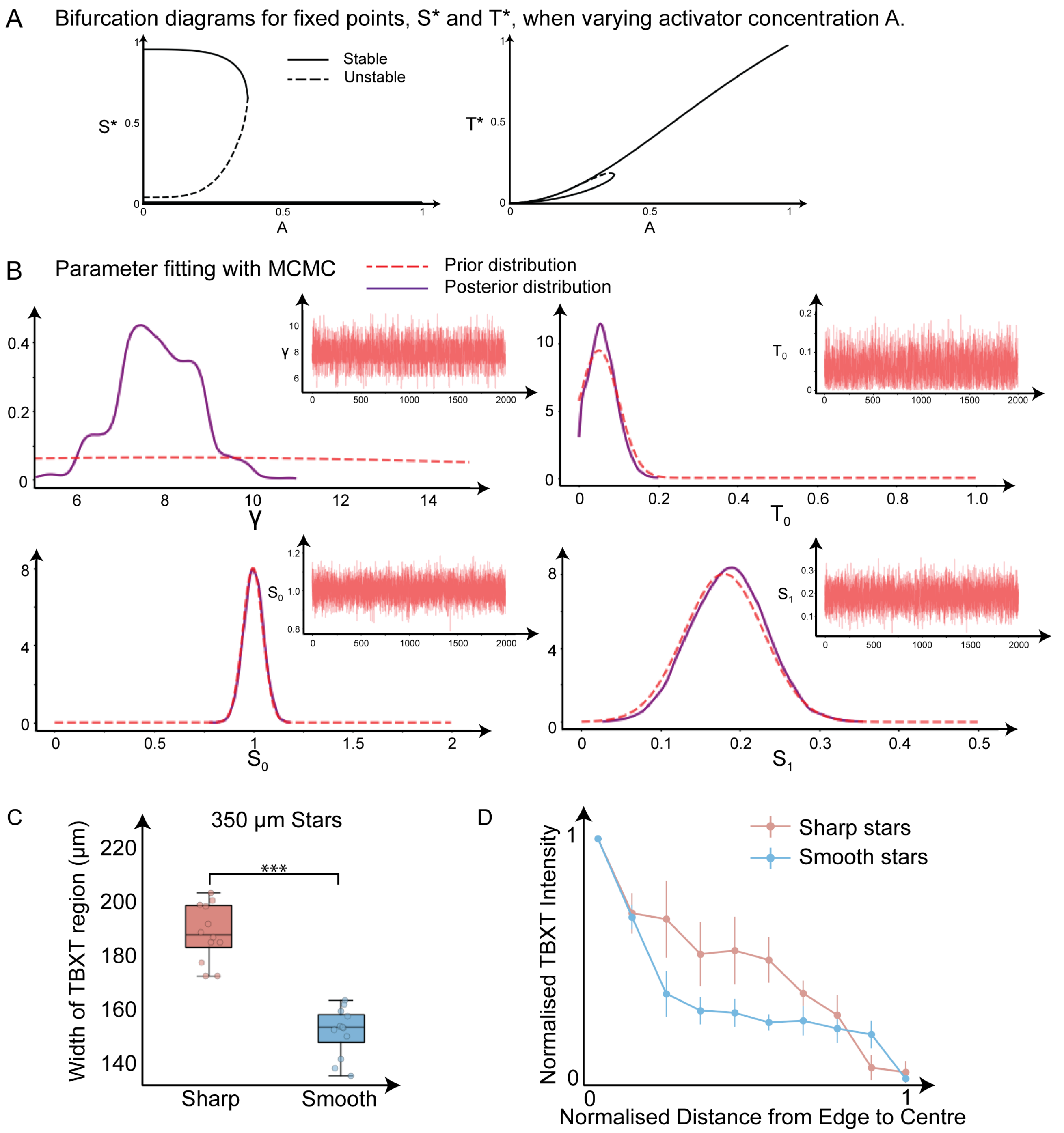


Figure S5: ***ODE system models SOX2 and TBXT dynamics*** **A.** Bifurcation plot when varying activator concentration $A$ for a set of parameters from Table 2. At low activator concentrations, the system is bistable but at high concentrations, there is only one stable fixed point. **B.** Parameter fitting using MCMC. Red dashed lines are manually chosen truncated normal distributions as prior distributions while purple solid lines are the posterior distributions after MCMC fit. The insets are MCMC trace plots. **C.** Widths of TBXT pattern at convex corners of sharp (pink) and smooth (blue) stars. There is a statistically significant difference between the widths (t-test, t=9.057, p-value$< 10^{-3}$). **D.** Normalised TBXT intensity profiles using the same method as in Fig. 3 on sharp and smooth stars.

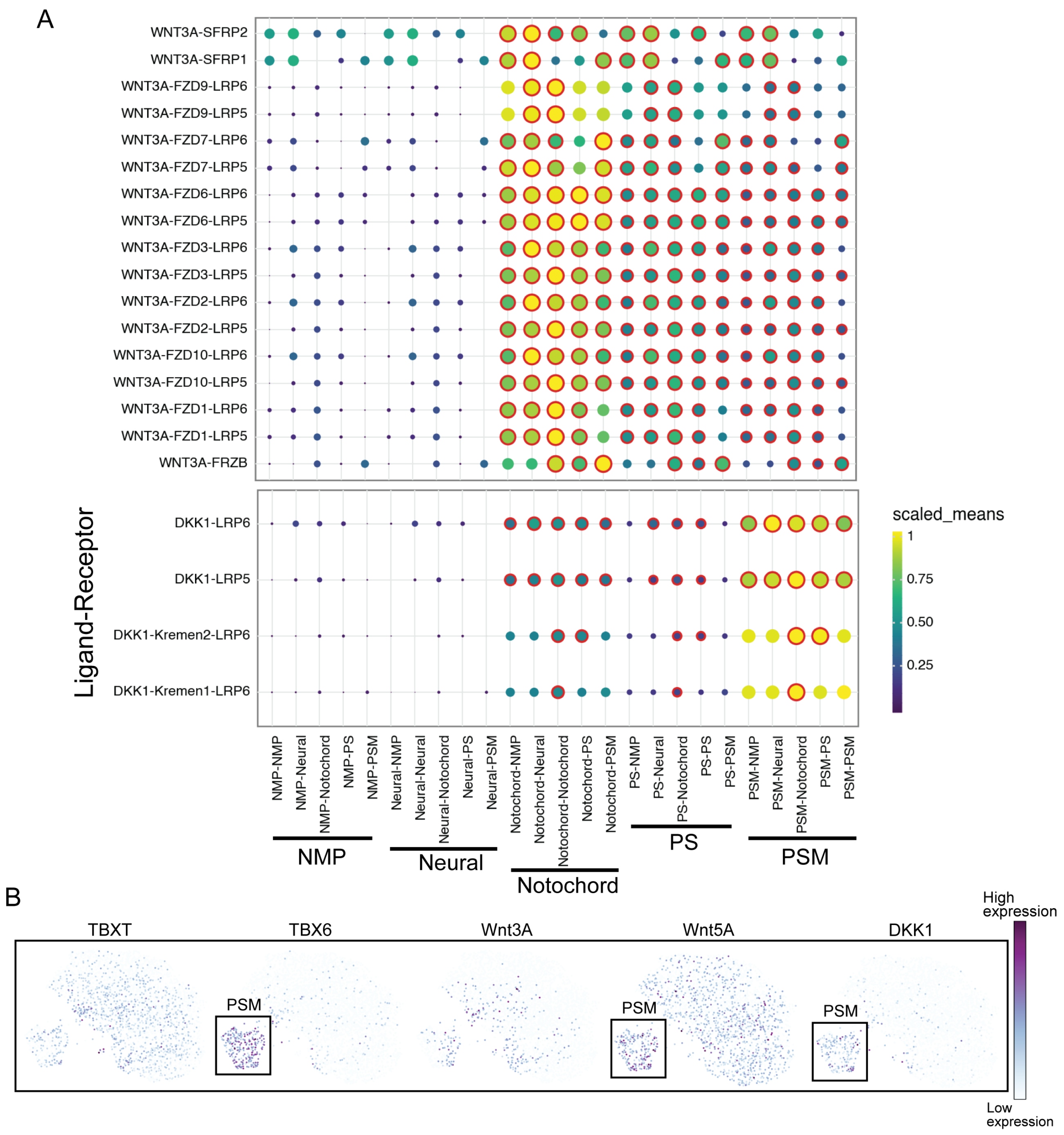


Figure S6: ***Analysis of scRNA-seq dataset of neuruloid at day 3 post-induction provided in[1] show presence of Wnt and Dkk ligands in PSM cell fates.*** **A.** Dot plots of activator-inhibitor pair, Wnt and Dkk interactions between different clusters. Dot colour and size represent the scaled mean expression of ligand–receptor pairs between cell types. Red outlines indicate statistically significant interactions (p¡0.05) based on CellPhoneDB permutation testing. **B.** Wnt and Dkk ligands are present in the system and are expressed in high levels in PSM, which are marked by high TBX6 expression. Since TBXT is required for commitment of NMPs to PSM, this indicates a good candidate of a potential activator-inhibitor pair driving the distinct patterning of TBXT profiles. *PS: primitive streak, PSM: presomitic mesoderm, NMP: neuromesodermal progenitors*

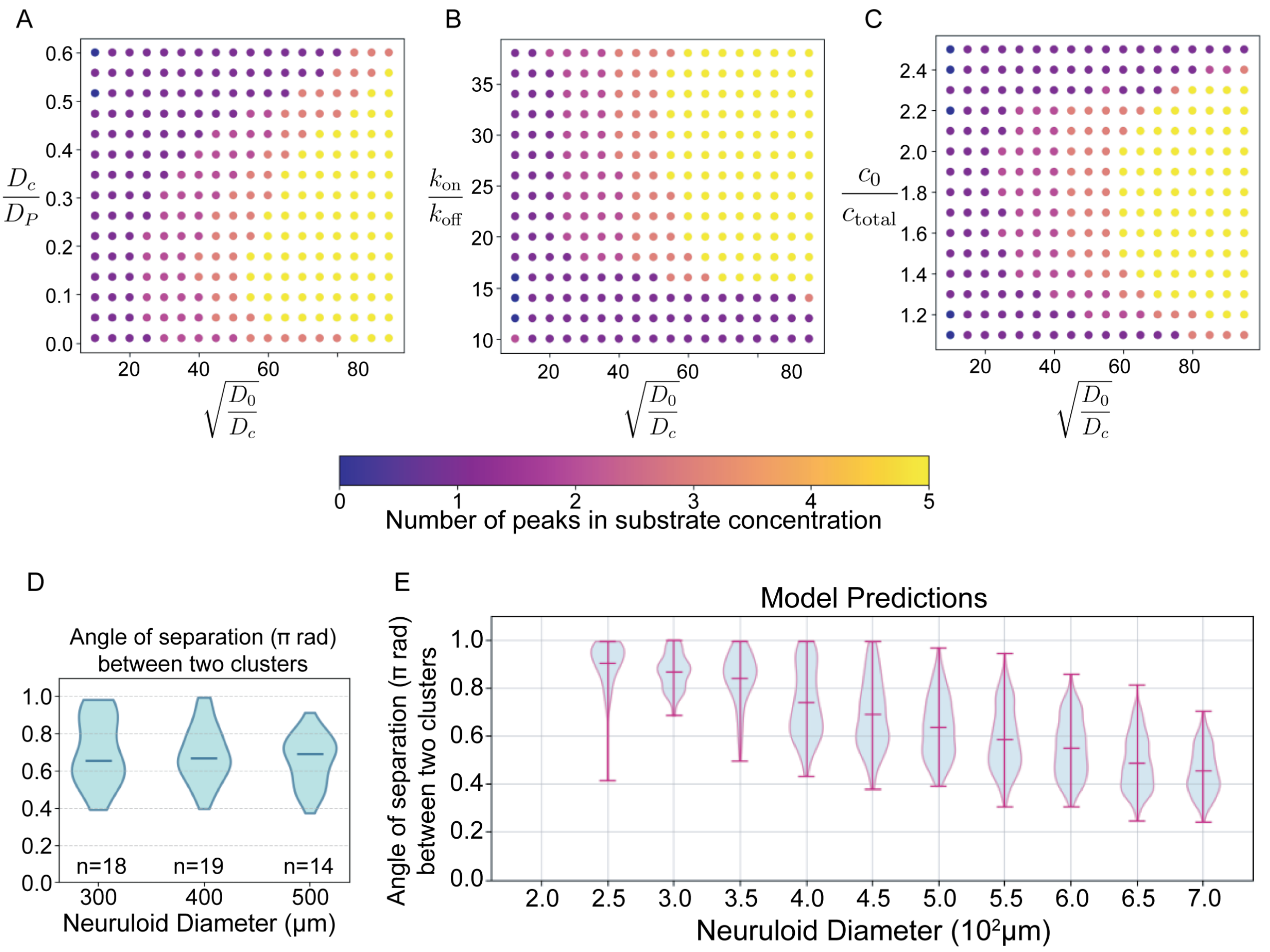


Figure S7: ***Substrate-depletion modelling predictions of cluster formation.*** **A-C.** Parameter search of the substrate-depletion model. The dimensionless parameter for x-axis in all plots is $\sqrt{D_0/D_c} = \sqrt{k_{\text{off}}L^2/D_c}$. Dimensionless parameter on the y-axes are **A.** $D_c/D_P$ **B.** $k_{\text{on}}/k_{\text{off}}$ and **C.** $c_0/c_{\text{total}}$, where $c_{\text{total}} = N/L$, $L$ is the length of the domain. Colours of the scatter plots represent the number of peaks in the substrate concentration when stable patterns are formed. If the parameter was not varied, then the fixed parameters are $D_c/D_P = 0.05, k_{\text{on}}/k_{\text{off}} = 20$ and $c_0/c_{\text{total}} = 1.5$. **D.** For higher resolution images, quantification of smallest separation angle between adjacent clusters in $\pi$ radians. **E.** Predictions on the separation angles between two clusters for increasing neuruloid diameter.
